# Acetylation of H3K115 is detected associated with fragile nucleosomes at CpG island promoters and active regulatory sites

**DOI:** 10.1101/2023.11.10.566531

**Authors:** Yatendra Kumar, Dipta Sengupta, Elias T. Friman, Robert S. Illingworth, Manon Soleil, Zheng Fan, Hua Wang, Kristian Helin, Matthieu Gerard, Wendy A. Bickmore

**Author notes:** **Correspondence to:** Y.K. or W.A.B: MRC Human Genetics Unit, IGC, Crewe Road, Edinburgh EH4 2XU, UK. These authors contributed equally (Yatendra Kumar, Dipta Sengupta).

## Abstract

Acetylation of lysine residues in the tail domain of histone H3 is well characterized, but lysine residues in the histone globular domain are also acetylated. Histone modifications in globular domain have regulatory potential because of their impact on nucleosome stability but remain poorly characterized. In this study we report the genome-wide distribution of acetylated H3 lysine 115 (H3K115ac), a residue on the lateral surface at the nucleosome dyad, using chromatin immunoprecipitation. In mouse embryonic stem cells, we find that detectable H3K115ac is enriched at the transcription start site of active CpG island promoters, but also at polycomb repressed promoters prior to their subsequent activation during differentiation. By contrast, at enhancers H3K115ac enrichment is dynamic, changing in line with gene activation and chromatin accessibility during differentiation. Most strikingly, we show that H3K115ac is detected as enriched on “fragile” nucleosomes within nucleosome depleted regions at promoters, and active enhancers where it coincides with transcription factor binding, and at CTCF bound sites. These unique features suggest that H3K115ac correlates with, and could contribute, to nucleosome destabilization and that it might be a valuable marker for identifying functionally important regulatory elements in mammalian genomes.

## Introduction

Post-translational modifications (PTMs) of histone tails play key roles in chromatin structure and gene regulation. Modifications of the histone tails have attracted attention due to the unstructured nature of the domain (Ghoneim et al., 2021; Peng et al., 2021) and accessibility to chromatin modifiers and readers. However, more recently modifications in the globular domains of core histones have been associated with various chromatin functions (Tropberger & Schneider, 2013; Pradeepa et al., 2016; Zorro Shahidian et al., 2021) but remain relatively understudied.

Modifications on the lateral surface of the histone octamer are of particular interest because these can directly affect interactions with DNA (Cosgrove et al., 2004; Lawrence et al., 2016). For example, acetylation of lysine-56 on histone H3 (H3K56ac) is predicted to increase accessibility at both the entry and exit sites of the nucleosome, impacting gene expression, DNA replication and repair (Rajagopalan et al., 2017; Rodriguez et al., 2019).

Lysine-64 on H3 is close to the nucleosome dyad axis and its acetylation (H3K64ac) decreases nucleosome stability (diCerbo et al., 2014). Lysine-122 of H3 (H3K122) is also located close to the dyad axis and its acetylation impacts nucleosome stability and weakens histone-DNA interactions (Manohar et al., 2009; Rajagopalan et al., 2017) with a direct consequence on transcription from a chromatinised template in vitro (Tropberger et al., 2013). H3K122ac also makes nucleosomes more susceptible to disassembly by chromatin remodellers (Chatterjee et al., 2015). Other acylation events, such as succinylation of H3K122 also impact nucleosome stability (Zorro Shahidian et al., 2021). Whilst H3.3 is not essential in mouse embryonic stem cells (mESCs), mutation of H3.3 K122 to alanine is cell lethal (Patty et al., 2024).

H3K115 is in the globular domain of H3 at the nucleosome dyad. Because both H3K115 copies in the histone octamer are juxtaposed on the lateral surface of the nucleosome in close contact with the overlying DNA (Figure 1A), acetylation of H3K115 has a high potential to disrupt histone-DNA interactions and influence nucleosome dynamics, nucleosome breathing and histone release (Zhou et al., 2019). Consistent with this, acetylation of H3K115 (H3K115ac) favours nucleosome disassembly and nucleosome sliding in vitro, by itself or together with H3K122ac (Manohar et al., 2009; Simon et al., 2011).

**Figure 1.**
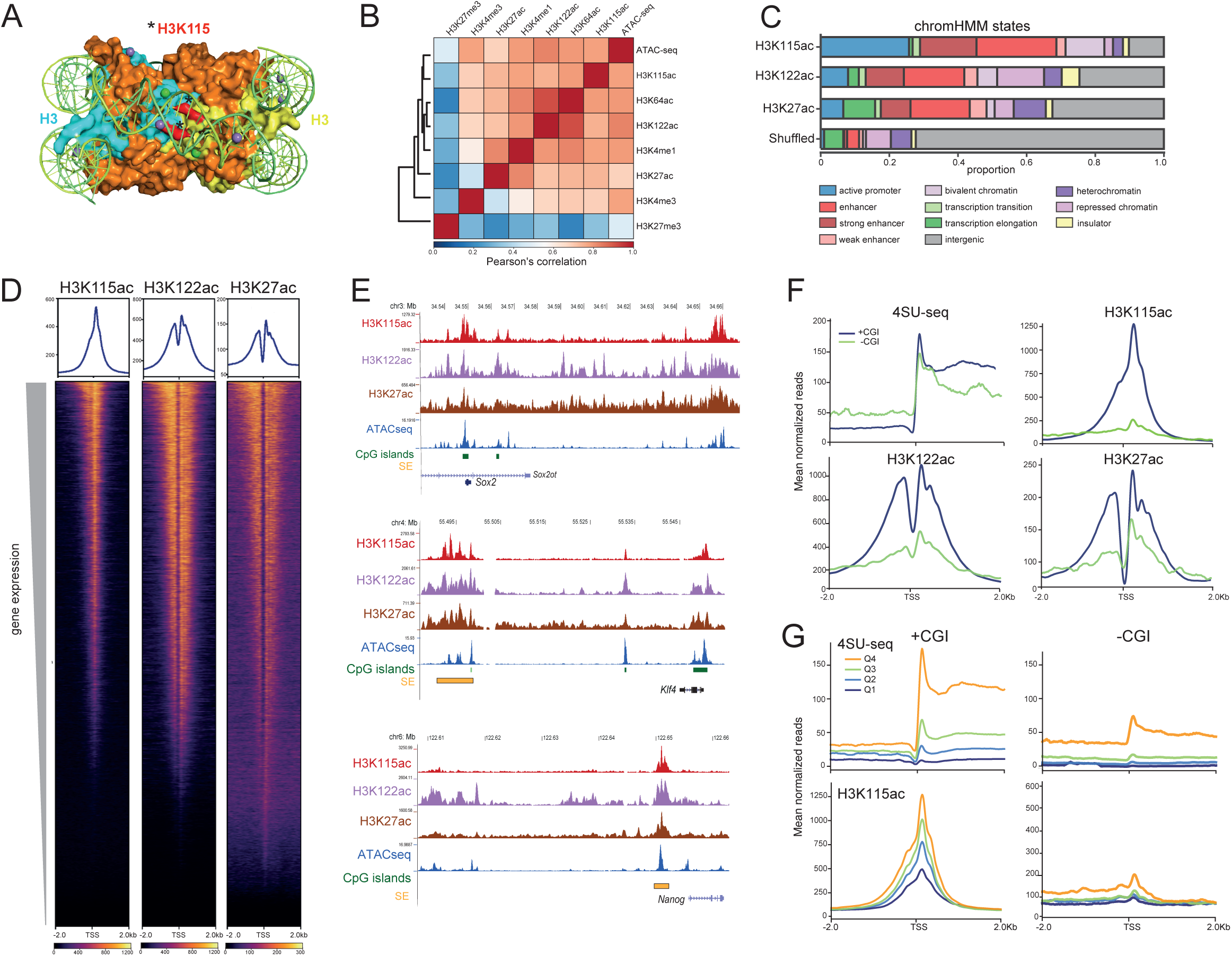
H3K115ac is associated with CGI promoters. A) Nucleosome structure looking down on the dyad axis (modified from PDB-5X7X, Taguchi et al., 2017). The two H3 molecules are shown in cyan and yellow, other histones are in orange and DNA in green. N-terminal histone tails are hidden. Both copies of H3K115 (red and asterisked) are juxtaposed at the dyad axis close to the overlying DNA. B) Pearson’s correlation matrix with hierarchical clustering in mESCs. Correlation is computed for read counts in 10 Kb windows across the genome for ATAC-seq data, ChIP-seq data for active (H3K122ac/H3K27ac/H3K27ac: GSE66023; H3K4me3: GSM1003756; H3K4me1: GSM1003750; and repressive (H3K27me3: GSM1276707), histone H3 modifications and for H3K115ac. C) Proportions of H3K115ac, H3K122ac and H3K27ac ChIP peaks that overlap genomic segments defined by chromHMM in the mouse genome (Ernst and Kellis, 2012; Pintacuda et al., 2017). D) Heatmap of H3K115ac ChIP-seq signal (this study), H3K122ac and H3K27ac (Pradeepa et al., 2016) in mESCs with respect to TSS of the top 50% of genes by expression and sorted by decreasing gene expression. E) UCSC genome browser screenshot showing ChIP-seq data for H3K115ac, H3K122ac, H3K27ac and ATAC-seq in mESCs at the *Sox2*, *Klf4* and *Nanog* loci. CpG islands (CGI) are indicated. Genome co-ordinates (Mb) are from the mm10 assembly of the mouse genome. F) Mean normalised reads of, (upper left) 4SU-seq centred at TSS of top 50% genes by expression (4SUseq tags in TSS +500bp region). Genes are divided into TSS that do (CGI+) or do not (CGI-) overlap with CpG islands. Upper right and lower panels show average profiles of H3K115ac, H2K122ac and H3K27ac ChIP-seq read-density at these same TSS classes. The higher H3K115ac read-density at CGI+ TSS is not due to sample size (Wilcox, p-value < 2.2e-16, normalized read coverage within a window spanning TSS +500bp). G) Average profiles (mean normalised reads) of, (top); 4SU-seq in mESCs centred at protein-coding TSSs (+/-2kb) divided into quartiles (Q1-Q4) of 4SU-seq signal within 500bp upstream and downstream of the TSS for (left) TSS overlapping a CGI (+CGI), or (right) promoters without any CGI (-CGI). Below; H3K115ac ChIP-seq signal around these TSS quartiles defined by 4SU.

Promoters have a specific nucleosome organization: two well-positioned nucleosomes (+1 and −1) flank a nucleosome depleted region (NDR) immediately upstream of the transcription start site (TSS). This view has been revised with the detection of micrococcal nuclease-(MNase) and salt-sensitive nucleosome species within the NDRs after limited chromatin digestion with MNase. These have been termed fragile, unstable, or partially unwrapped nucleosomes (Henikoff et al., 2011; Kent et al., 2011). Fragile nucleosomes can be isolated by MNase digestion under mild conditions and contain the H3.3 and H2AZ histone variants (Jin and Felsenfeld, 2007). In mouse embryonic stem cells (mESCs), chemical cleavage methods have also identified sub-nucleosomes at TSSs (Ishi et al., 2015; Voong et al., 2016). To the best of our knowledge, fragile nucleosomes have not yet been associated with a specific histone modification in mammalian cells.

H3K64ac and H3K122ac have been detected by chromatin immunoprecipitation (ChIP) at the TSS of active genes and at some active enhancers in mammalian cells (Tropberger et al., 2013; diCerbo et al., 2014; Pradeepa et al., 2016). In this study we report the distribution of H3K115ac, as detected by ChIP, across the mammalian genome in mESCs, and during the differentiation to neural progenitor cells (NPCs). This reveals the focal association of H3K115ac with fragile nucleosomes at both active and repressed CpG island (CGI) promoters, at active enhancers, and at other sites important for genome architecture. Our data suggest that H3K115ac ChIP-signal has a genomic distribution distinct from other H3 acetylation marks, and that it could be an important new tool for the functional annotation and understanding of mammalian genomes.

## Results

### H3K115ac is associated with active regions with a strong preference for CpG island promoters

To determine where H3K115 acetylated nucleosomes can be detected across the mammalian genome, we used a commercially available antibody and confirmed its specificity for H3K115ac and not for unmodified H3K115, H3K115R, or H3K122ac by dot blotting against synthetic peptides (Figure S1A) and for a variety of H3 and H4 lysine post-translational modifications by chromatin immunoprecipitation ChIP of bar-coded nucleosomes (Figure S1B).

Using this antibody, we then performed native ChIP followed by next generation sequencing (ChIP-seq) from mESCs. H3K115ac ChIP data were highly reproducible between biological replicates (Pearson’s correlation *r*=0.97, Supplementary table 1). Genome-wide analysis showed that H3K115ac ChIP peaks correlate with regions decorated with histone H3 post-translational modifications (PTMs) associated with active gene regulation, both on the H3 tail (H3K27ac, H3K4me1) and at the lateral surface of the nucleosome (H3K64ac and H3K122ac) as well as with regions of chromatin accessibility as assayed by the ATAC-seq data we generated (Pearson’s correlation *r*=0.97 between replicates, Supplementary table 1) (Figure 1B). Using a chromHMM segmentation approach, we observed that H3K115ac regions have a greater degree of overlap with active promoters and enhancers, compared to H3K122ac and H3K27ac (Figure 1C). Notably, there are fewer called peaks for H3K115ac (∼39000), compared to other H3 lysine acetylation marks in mESC (H3K122ac ∼80,000; H3K64ac ∼65,000; H3K27ac >100,000).

Approximately one third of detected H3K115ac peaks are associated with transcription start sites (TSS), and heat maps show H3K115ac levels generally increasing with the transcription levels as assayed by 4SU-seq (Benabdallah et al., 2019) (Figure 1D). However, while examining key pluripotency genes, we noticed that H3K115ac enrichment, along with that for H3K64ac and H3K122ac, is detected at the TSS of *Sox2* and *Klf4*, but not that of *Nanog*, despite an ATAC-peak being detected at the Nanog TSS indicating that H5K115ac detection is not simply a feature of all open chromatin (Figure 1E). The *Sox2* and *Klf4* promoters are located within CpG islands (CGIs), but the promoter of *Nanog* lacks a CGI, prompting us to investigate H3K115ac with respect to CGIs. Genome wide, ∼60% of CGI associated TSSs (CGI promoters) are detected as associated with H3K115ac in mESCs compared to just 2% for non-CGI promoters. Plotting mean normalised reads around (+/-2kb) of TSS shows that CGI promoters exhibit H3K115ac levels many fold higher than non-CGI promoters with comparable levels of transcription (Figure 1F). This same discrimination between CGI and non-CGI promoters is not so pronounced for H3K122ac and H3K27ac ChIP data.

Non-CGI promoters have lower overall levels of transcription compared to CGI promoters, and for this promoter class H3K115ac enrichment detected by ChIP is only really seen for the highest quartile of transcription (4SU) quartile of expression (Figure 1G). CGI promoters on the other hand, exhibit significant levels of detected H3K115ac even for the lowest quartile of expression. These results suggest a special link between CGI promoters and H3K115ac.

### H3K115ac is detected at CpG island promoters poised for activation

To determine the relationship between the detection of H3K115ac and dynamic gene expression, we differentiated 46c mESC cells to neural progenitor cells (NPCs) monitoring differentiation by Sox1-GFP fluorescence. ChIP-seq for H3K115ac (Pearson’s correlation *r*=0.90, Supplementary table 1) and ATAC-seq (Pearson’s correlation *r*=0.94, Supplementary table 1) were carried out after 7 days of differentiation and nascent RNA quantified (4SU-seq) on days 0, 3, 5 and 7 (Benabdallah et al., 2019).

TSSs were classified into those that gained or lost H3K115ac ChIP signal during differentiation if the TSS (−500 to +500 bp) showed at least a 2-fold change in H3K115ac signal compared with day 0. Average levels of transcription increased from TSS that also gained H3K115ac, and there was loss of H3K115ac from downregulated TSS (Figure 2A).

**Figure 2.**
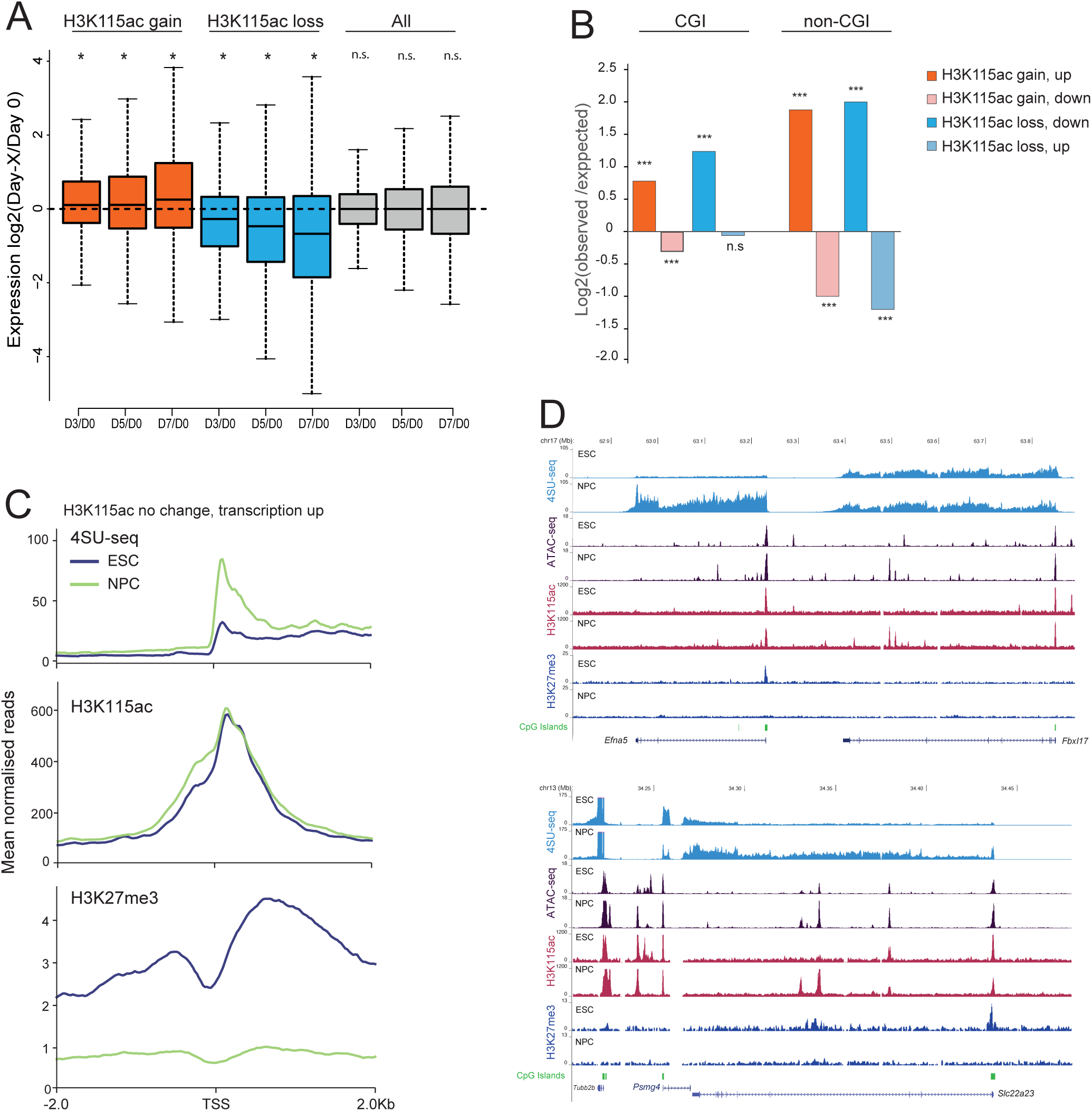
Changes in H3K115ac during differentiation. A) Boxplots displaying the changes in activity (4SU-seq) of promoters that gain or lose H3K115ac ChIP signal across the 7 days of mESC to NPC differentiation. Log2 fold change is shown relative to day 0. *Paired Wilcox, p <0.01. B) Bar plot showing enrichment of gene sets defined based on differential H3K115ac (gain/loss) and differential expression during differentiation(up/down). Enrichments are calculated for CGI and non-CGI promoters. *** Fisher’s exact test p < 0.01; n.s – p > 0.01, supplementary table 2. C) Aggregate profile plots for 4SU-seq, and ChIP-seq data for H3K115ac and H3K27me3 (Mikkelsen et al., 2007) from mESCs and NPCs at promoters with no significant change in H3K115ac occupancy, but significant transcriptional upregulation during differentiation. D) UCSC genome browser screenshot showing 4SU-seq, ATAC-seq, and ChIP-seq data for H3K115ac, and H3K27me3 (Mikkelsen et al., 2007) in mESCs and differentiated NPCs at the *Efna5*, and *Slc22a23* loci. CpG islands (CGI) are indicated. Genome co-ordinates (Mb) are from the mm10 assembly of the mouse genome.

Given the strong bias of H3K115ac enrichment toward CGI promoters largely independent of transcription (Figure 1F), we classified CGI- and non-CGI promoters based on changes in H3K115ac (gain/loss) and expression (up/down) during differentiation (Figure 2B).

Promoters exhibit positive enrichment (log_2_observed/expected) for congruent changes of H3K115ac and transcription (gain ∼ up and loss ∼ down) and negative enrichment for incongruent changes (gain ∼ down and loss ∼ up) (Figure 2B, supplementary table 2).

However, these enrichments are stronger for non-CGI promoters than for CGI promoters. This is consistent with our observation that some CGI promoters have significant levels of H3K115ac detectable while being transcribed at extremely low levels (Figure 1G).

In line with this, we identified a set of promoters that are transcriptionally upregulated in NPC but without any significant changes in the levels of H3K115ac detected (Figure 2C). In mESC, these promoters exhibit very low transcription along with high levels of H3K27me3, which are then lost upon differentiation (Goronzy et. al., 2022). Compared to these promoters, a background set of promoters with no detected changes in H3K115ac and no change in transcription (> 2 fold) during differentiation have very low levels of H3K27me3 (Figure S2A). H3K115ac-marked promoter sets - including those showing no detectable change in H3K115ac despite transcriptional activation during differentiation - are enriched for high-confidence bivalent genes taken from another study (Seneviratne et. al., 2024, supplementary table 2). This indicates that H3K115ac, as assayed by ChIP, is a feature of CGI associated genes that are polycomb targets in mESCs and that are poised for later activation in NPCs (Figure S2B). Examples of this include *Efna5*, *Slca23* (Figure 2D) and *Insm1* (Figure S2C), all genes known to have a function in neuronal cells.

H3K115ac shows a stronger enrichment at CGI, polycomb repressed and bivalent promoters in mESC than two other globular domain H3 modifications - H3K64ac and H3K122ac (Figure S2D). H3K27ac, as expected, is depleted from CGI promoters, since H3K27ac and H3K27me3 are mutually exclusive. In contrast to H3K115ac, H3K64ac and H3K122ac are enriched over the gene body and upstream regions, but not at the TSS, of CGI-associated genes (Figure S2E).

### H3K115ac is associated with fragile nucleosomes within promoter NDRs

We noted that, H3K115ac ChIP signal over promoters appears much more focal and centred over the TSS compared with both H3K27ac and H3K122ac that are depleted over the TSS (Figure 1D). This suggests that H3K115ac may be associated with MNase-sensitive nucleosomes in the promoter NDR. These nucleosomes can be detected as sub-nucleosomes (footprint < 147bp) obtained from small fragments sequenced from partial MNase digestion (Carone et al., 2014) or mono-nucleosomes (footprint ∼ 147 bp) obtained under low salt conditions (Jin & Felsenfeld, 2007). Both species of nucleosome particles are referred to as fragile nucleosomes (Carone et al., 2014; Ishi et al., 2015; Voong et al., 2016).

To investigate the distribution of H3K115ac in relation to nucleosome size, we repeated the H3K115ac and H3K27ac ChIP on partially MNase-digested native chromatin from mESCs and sequenced both ends of the purified DNA fragments (paired-end) (Pearson’s correlation *r*=0.99 between biological replicates for both H3K115ac and H3K27ac, Supplementary table 1). Around (+/- 1kb) the most active promoters (4SU Q4), mean signal from small (<150 bp) MNase digested fragments accumulate inside the TSS NDR (Figure S3A). Signal from fragments > 150 bp exhibits the characteristic nucleosome depletion, with the NDR becoming more pronounced with increasing fragment lengths. We used this rationale to split ChIP fragments into sub-nucleosomes (≤150 bp) and mono-nucleosomes (151 – 230 bp) (Figure S3B). Paired-end ChIP-seq confirmed H3K115ac detection within the NDR of active promoters on sub-nucleosomes and mono-nucleosomes immediately upstream of the TSS (Figure S3C). In contrast, H3K27ac signal is enriched in the gene body and upstream of the TSS but is depleted over the NDR.

H3K115ac ChIP preferentially selects nucleosomes that are shorter than the average input nucleosomes. This is not the case for nucleosomes selected by H3K27ac ChIP (Figure S3D, Supplementary table 3). This is an indication that H3K115ac may be associated with destabilized and partially unwrapped fragile nucleosomes. Because nucleosomes associated with A/T-rich sequences are known to be more sensitive to MNase compared to sites of higher G/C content (Chereji et al., 2019), we analysed A/T content in H3K115ac and H3K27ac libraries. Sequences associated with H3K115ac sub- and mono-nucleosomes have significantly higher A/T content than those marked with H3K27ac (Figure S3E, p < 0.01, Wilcox Test, supplementary table 3). This bias does not originate from differences in MNase digestion levels, because the matched input libraries do not show such a difference. This suggests a stronger association of H3K115ac detection with nucleosome fragility compared to of H3K27ac. Difference in A/T content also suggests that H3K115ac and H3K27ac fragments come from distinct populations of nucleosomes. H3K115ac modified sub-nucleosomes and mono-nucleosomes have the same A/T content, indicating that H3K115ac nucleosomes are present in different stages of unwrapping or stability (Figure S3E).

To explore H3K115ac ChIP paired end fragment length(bp) from mESCs in relation to promoter NDRs and flanking nucleosomes, we plotted fragment length against the distance between the ChIP fragment centre and the nearest TSS as 2D contour V-plots of highest density regions for a conservative TSSs set, with colour coding of the plots showing the proportion of ChIP fragments within a particular contour (see Methods). Signal from input libraries show the promoter NDR flanked by positioned nucleosomes with smaller nucleosome particles of varying sizes inside the NDR (Figure 3A). H3K27ac signal is most abundant on the +1 nucleosome with barely detectable signal inside the NDR. In contrast, H3K115ac is most enriched on the −1 nucleosome and on subnucleosome-sized fragments inside the NDR (Figure 3A, bottom panel). We then defined NDRs by taking the distance between the TSS and the −1 nucleosome positioning data from published datasets (West et al., 2014; Voong et al., 2016), scaled them to the same length, and plotted the coverage of stable nucleosomes (MNase-seq), H3K27ac and H3K115ac fragments at these sites (Figure 3B). H3K27ac is generally depleted from the NDR, but both H3K115ac modified mono- and subnucleosomes are enriched within the NDR. Within the NDR, subnucleosomes-sized H3K115ac MNase fragments are TSS proximal whereas mononucleosomal fragments are displaced toward the −1 nucleosome (Figure 3B).

**Figure 3.**
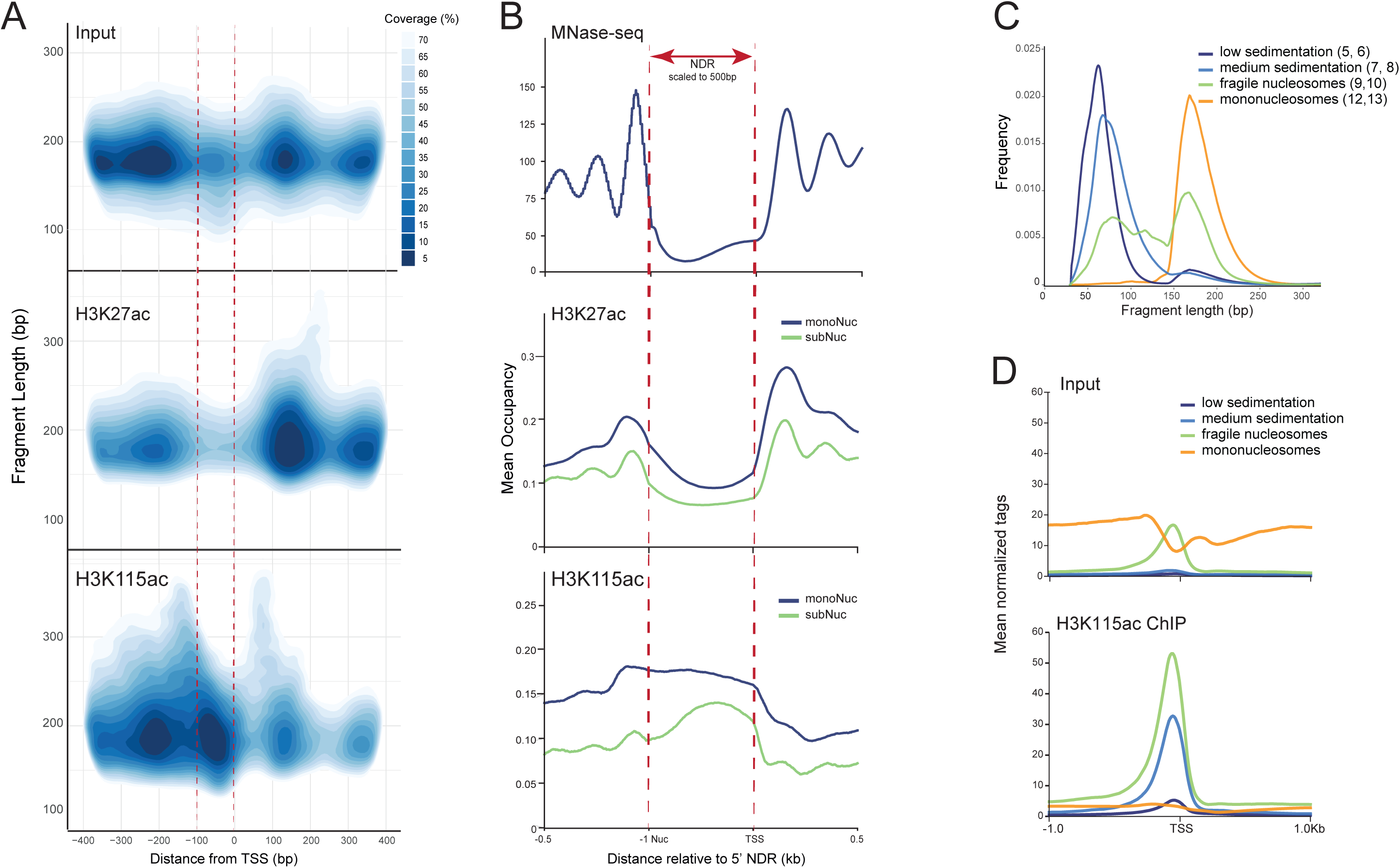
H3K115ac is associated with fragile nucleosomes. A) Contour plots depicting the high-density regions of chromatin fragments around TSS (+/-400bp) as a function of fragment length (bp) generated from; (top) Input MNase library, (centre) H3K27ac ChlP and (bottom) H3K115ac ChIP. Coverage refers to the proportion of ChIP-fragments in the indicated color-coded contours. The region from 100 bp upstream to the TSS is indicated with dashed lines in red. B) Mean nucleosome occupancy around mouse TSSs plotted with respect to the NDRs, scaled to a length of 500bp. MNase-seq data (top, West et al., 2014) were used to define NDR as the region between the 3’ boundary of-1 nucleosome and the TSS. Mean occupancy of (middle) H3K27ac or (bottom) H3K115ac ChlP-seq fragments is split into sub- and mono-nucleosomes around the scaled NDRs. C) Fragment length distribution (bp) of MNase-digested native chromatin fractionated with sucrose gradient sedimentation. Fractions with different nucleosome species (based on the fragment length) were pooled (indicated in parentheses). D) Input (top) and H3K115ac ChIP-seq (bottom) data, centred at TSS (+/- 1kb) of most active genes (top 25%) in mESCs, performed on different nucleosome species isolated with sucrose gradient sedimentation from panel C. Data is spike-in normalized.

To further investigate the properties of subnucleosomal and mononucleosomal particles, we separated MNase digested mESCs chromatin by sucrose gradient sedimentation which can separate fragile and stable nucleosomes (Nocente et al., 2024) (Figure 3C). Sequencing of the chromatin fractions showed the expected depletion of mononucleosomes at the TSS NDR, whereas the slower sedimenting fragile nucleosomes are enriched over the NDR (Figure 3D). H3K115ac ChIP-seq from chromatin fractions showed that H3K115ac is detected as enriched in fragile nucleosomal and even slower sedimenting (lower mass) subnucleosomal fractions at TSS, but not in fractions with stable mononucleosomes (Figure 3D).

### H3K115ac modified nucleosomes overlap transcription factor binding sites at active enhancers

With chromHMM segmentation, the largest proportion of H3K115ac ChIP peaks overlap regions identified as putative enhancer elements (Figure 1C). Indeed, H3K115ac peaks decorate the known super enhancers of *Sox2*, *Klf4* and *Nanog*, even though the latter gene lacks H3K115ac at its non-CGI promoter (Figure 1E). Peaks detected by ChIP for H3K115ac overlap with a subset of enhancers that are marked by H3K27ac and H3K122ac and that exhibit the highest chromatin accessibility as assayed by ATAC-seq compared to enhancers marked with H3K27ac and H3K122ac, alone and in combination (Figure 4A).

**Figure 4.**
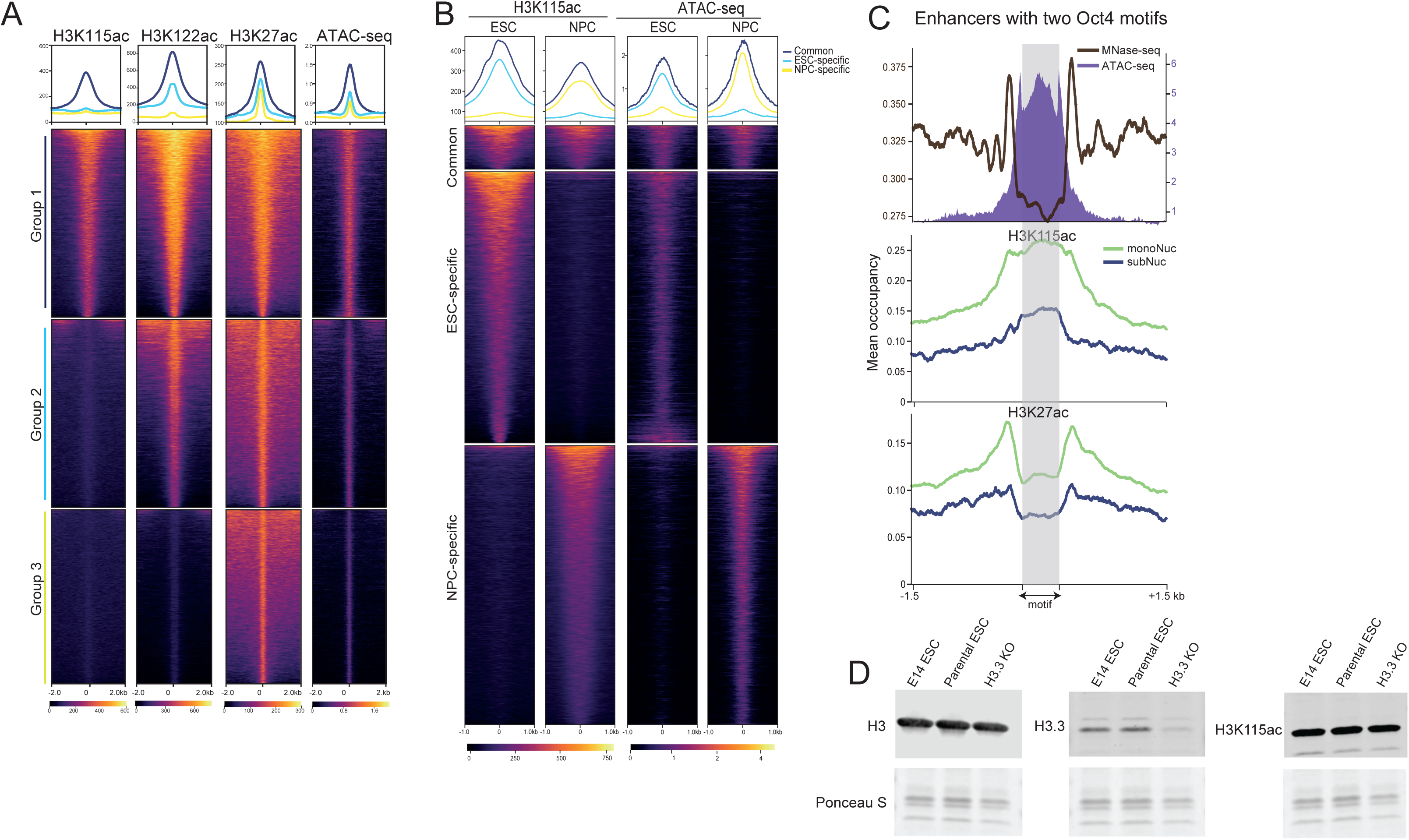
H3K115ac marks active enhancers. A) Heatmap showing the coverage of H3K115ac, H3K122ac, H3K27ac and ATAC-seq centred on promoter distal accessible peaks (putative enhancers) in mESCs. Data is grouped in enhancers marked by all three H3 acetylation marks (Group 1; H3K27ac+ H3K122ac+ H3K115ac+), just H3K27ac together with H3K122ac (Group 2), or H3K27ac alone (Group 3). B) Heatmaps showing H3K115ac and ATAC-seq signal for common and dynamic enhancers between ESC and NPC. Loss/gain of H3K115ac correlates with loss/gain in chromatin accessibility. C) mESC enhancers selected based on the presence of two Oct 4 motifs (n=650) within the Tn5 accessible region, with the region between the two motifs scaled to the same length (shaded grey region). Top; MNase-seq signal (black, left y axis) and ATAC-seq (purple, right y axis). H3K115ac ChlP-seq (middle panel) and H3K27ac ChlP-seq (bottom panel) on mononucleosomes (monoNuc; green) or subnucleosome-sized fragments (subMuc; blue). D) Immunoblotting for H3, H3.3 and H3K115ac on whole cell extracts from E14 mESCs, H3.3 knock-out ESCs and the parental ESC line. Ponceau S staining of histones shown below as loading control. Full gel with H3, H3.3 and H3K115ac (green) and H4 (red) LiCor staining, and a biological replicate blot are shown in Fig S4H.

Sequences capable of enhancer activity on episomes in mESCs have been identified by STARR-seq (Peng et al., 2020) but only half of these correspond to accessible endogenous chromatin sites, as determined by ATAC-seq. Compared with H3K27ac and various acetyl-lysines on histone H2B (Narita et al., 2023), H3K115ac detected by ChIP better discriminates those STARR-seq hits mapping to accessible (ATAC+) sites in the mESC genome from those that are ATAC- (Figure S4A), even though the proportions of total H3K115ac and H2B acetylated lysine peaks in the mESC genome that are ATAC+ are similar (Figure S4B). Association of H3K115ac with active enhancers is further substantiated by the enrichment of transcription factors, co-activators and RNA polymerase II at non TSS H3K115ac regions as revealed by a random forest approach using published datasets (Figure S4C).

During mESC to NPC differentiation, the most frequent changes in H3K115ac detection occur at enhancers (Figure S4D). Loss of H3K115ac signal from ESC-specific enhancers correlates to loss of chromatin accessibility (ATAC-seq), enhancers gaining H3K115ac signal during differentiation to NPCs also gained chromatin accessibility (Figure 4B).

Like promoters, active enhancers are known to be depleted of stable nucleosomes (Oruba et al., 2020. We therefore analysed H3K115ac and H3K27ac paired-end ChIP-seq data at mESCs enhancers occupied by the pluripotency TF Oct4 (King and Klose, 2017; MacCarthy et al., 2022). As expected, chromatin accessibility, as assayed by ATAC-seq, scales with Oct4 occupancy, with a sharp dip in accessibility centred at the Oct4 motif likely due to the protection offered by Oct4 binding (Figure S4E). H3K27ac levels correlate with Oct4 occupancy in regions immediately flanking the Oct4 binding site but are depleted over the binding site itself (Figure S4F). H3K115ac ChIP profiles also correlate with Oct4 occupancy, but both mononucleosomal and subnucleosomal H3K115ac marked fragments are enriched over the Oct4 motif itself (Figure S4F).

As Oct4 binds co-operatively (Sinha et al., 2023), we also analysed enhancers with two Oct4 motifs scaling the distance between the two motifs to a window of the same width. The presence of two Oct4 motifs in proximity creates sharp protected footprints in ATAC-seq data, with high chromatin accessibility between the two sites (Figure 4C, top panel). The absence of stable nucleosomes between the two motifs is evident in MNase-seq data, with strongly positioned nucleosomes flanking the ATAC-accessible region. H3K27ac is enriched on these flanking nucleosomes but is depleted between the Oct4 motifs (Figure 4C, bottom panel). In contrast, enrichment of H3K115ac ChIP signal on both mono- and sub-nucleosome-sized fragments spans the entire region between the two Oct4 motifs (Figure 4C, middle panel).

Active enhancers and promoters are known to harbour fragile nucleosomes carrying the histone variant H3.3 (Jin and Felsenfeld, 2007), with the −1 nucleosome reported to be hyperdynamic because of high H3.3 turnover (Schlesinger et al., 2017). Consistent with H3K115ac enrichment on the −1 nucleosome (Figure 3A), H3K115ac-marked distal elements correlate with sites of high H3.3 turnover (Deaton et al., 2016) (Figure S4G). To investigate if H3K115ac then merely correlates with fragile nucleosomes because it is deposited solely on H3.3, we generated a histone H3.3 knock-out mESC line. Immunoblotting shows that global levels of H3K115ac are not affected by H3.3 deletion indicating that H3K115ac is not specific to H3.3 (Figure 4D and Figure S4H). These data show that H3K115ac and its association with fragile nucleosomes, is linked to binding of TF complexes and chromatin accessibility at active enhancers elements and CGI promoters in an H3.3-independent manner.

### H3K115ac is detected at fragile nucleosomes at CTCF binding sites

Though CTCF binding was originally thought to exclude nucleosomes, fragile nucleosomes and subnucleosomal fragments have been detected at CTCF binding sites (Voong et al., 2016; Klein et al., 2023). MNase-seq data shows two arrays of well positioned stable nucleosomes on both sides of the CTCF motif, correlated with the level of CTCF occupancy, and consistent with the known ability of CTCF to create arrays of well positioned nucleosomes (Fu et al., 2008; Owens et al., 2019) (Figure 5A). H3K27ac is enriched on these +1 and −1 mononucleosomes, levels are low on subnucleosomal-sized fragments (Figure S5A) and H3K27ac is depleted from the CTCF NDR (Figure S5A). H3K1115ac levels also correlate with CTCF occupancy. H3K115ac modified mononucleosomes are enriched at the −1 and +1 positions relative to the CTCF motif, but strikingly H3K115ac modified subnucleosomal particles are enriched within the CTCF NDR and overlap the CTCF motif (Figure 5B).

**Figure 5.**
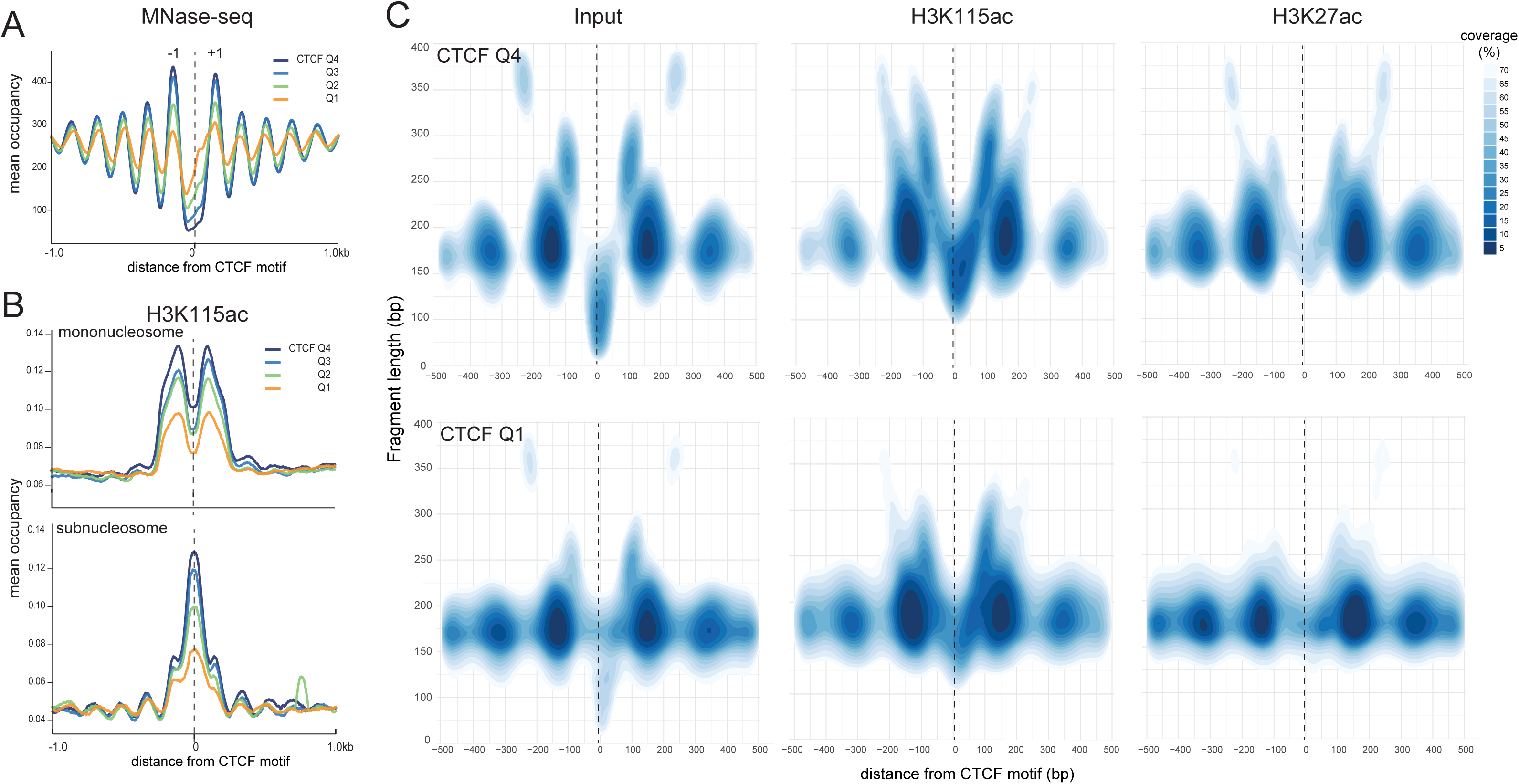
H3K115ac marks fragile nucleosomes at sites of high CTCF occupancy. A) Mean occupancy of nucleosomes derived from MNase-seq (from West et. al., 2014) around CTCF sites across the four quartiles of CTCF ChlP-seq peak strength. All CTCF motifs are oriented from 5’ to 3’ (left to right). Positions of first flanking upstream (−1) and downstream nucleosome (−1) positions are marked. B) H3K115ac ChlP-seq signal from (top) mononucleosomal and (bottom) subnucleosomal sized fragments in mESCs around CTCF motifs across the four quartiles of CTCF ChlP-seq peak strength as in (A). C) Contour plots depicting high density regions of (left) input, (centre) H3K115ac ChlP, (right) H3K27ac ChlP paired-end sequenced MNase fragments around top (Q4) and bottom (Q1) quartiles of CTCF ChlP-seq peaks in mESCs (data from Mas et al., 2018) as a function of fragment length. All CTCF motifs are oriented in the same direction.

Using contoured V-plots, we detected small (∼50-150 bp) and 250bp MNase resistant fragments within the CTCF NDR, flanked by strongly positioned nucleosome-sized fragments (compare Q4 to Q1 in Figure 5C, left panels). In agreement with average occupancy plots (Figure 5B), H5K115ac is detected on the −1 and +1 positioned nucleosomes and these nucleosomes exhibit a wider range of fragment sizes (Figure 5C, middle). This suggests that these nucleosomes are more dynamic than the stable nucleosomes visible in MNase-seq (Figure 5A). Within the CTCF NDR, H3K115ac ChIP fragments produce a V-shape pattern of which is more prominent at high CTCF occupancy sites (Figure 5C, middle). In the centre of the V, 100-175 bp sized fragments most likely contain H3K115ac marked subnucleosomes in complex with CTCF. The horns of this V-shaped pattern indicate H3K115ac-marked flanking mononucleosomes interacting with CTCF and/or other factors leading to MNase protected 225-350 bp fragments. In contrast, H3K27ac is depleted from the NDR, and its enrichment is largely limited to mononucleosomes immediately flanking the CTCF site (Figure 5C, right).

V-plots also reveal a subtle asymmetry in the distribution H3K115ac fragments within the CTCF NDR, with occupancy of subnucleosomal H3K115ac fragments slightly higher toward the 3’ end of the CTCF motif. While upstream nucleosomes (MNase-seq) maintain phasing, with decreasing CTCF occupancy the +1 nucleosome becomes fuzzier and begins to encroach into the CTCF NDR at low CTCF occupancy (Figure S5B). This directional bias is lost when the orientation of CTCF motifs is randomised. To ascertain that the directional bias is not due to higher CTCF occupancy merely coinciding with TAD boundaries, we used proximity to the nearest TAD boundary (Bonev et al, 2017) as the probability with which a given CTCF site participates in TAD boundary formation, and divided CTCF motifs into quartiles of this distance (TAD score; Supplementary table 4). We plotted occupancy of stable nucleosomes around the motifs in these quartiles and sorted each quartile by descending order of CTCF occupancy (Figure S5C). The fuzziness of nucleosomes downstream of CTCF sites is inversely correlated with CTCF occupancy across all TAD scores quartiles.

## Discussion

Lysine acetylation in the globular domain of histones, and particularly at the lateral surface of the nucleosome where histones are in close contact with overlying DNA, is not well studied but is important for nucleosome dynamics, nucleosome stability (diCerbo et al., 2014; Chatterjee et al., 2015; Lawrence et al., 2016; Rajagopalan et al., 2017) and transcription (Tropberger et al., 2013). In this study we explored acetylation of H3K115, a residue located at the nucleosome dyad which weakens histone DNA interactions and destabilizes nucleosome in vitro (Chatterjee et al., 2015). In *Saccharomyces cerevisiae* H3K115A and H3K115Q abrogate transcriptional silencing and result in hypersensitivity to DNA damage (Hyland et al., 2005). In *Drosophila melanogaster*, both H3K115R and H3K115Q mutations are embryonic lethality (Graves et al., 2016) indicating the biological importance of this H3 residue.

We found that in mESCs, and their differentiated derivatives, H3K115ac exhibits a strong preference for active and epigenetically repressed CGI promoters. This is consistent with the known correlation between CGIs and nucleosome depletion independent of transcription (Tazi and Bird, 1990; Fenouil et al., 2012), their enrichment in the variant histone H2A.Z, and the presence of specific CGI binding proteins that open chromatin (Yukawa et al., 2014; Grand et al., 2021). As well as active CGI promoters, H3K115ac is associated with polycomb repressed promoters in mESCs. Though polycomb complexes can be found at “bivalent” chromatin domains, characterized by the simultaneous presence of both repressive (H3K27me3) and active (H3K4me3) histone marks at the same promoter regions (Voigt et al., 2013), to our knowledge, H3K115ac is the first histone acetylation found within the NDRs associated with polycomb repressed promoters.

Histone acetylation, particularly in the histone H3 tail, has been used as a marker to identify active enhancers. H3K27ac is most widely used for this purpose (Moore et al., 2020; Stunnenberg et al., 2016) and various acetyl-lysines in the N-terminal of histone H2B (H2B-NTac) enhance the discovery of active enhancers when used in combination with H3K27ac (Peng et al., 2020). However, H3K27ac is dispensable for gene activation in mESCs during the exit from pluripotency and for enhancer chromatin accessibility (Zhang et al., 2020, Sankar et al., 2022) and it performs poorly as a predictor of functional variants in enhancers (Biddie et al., 2024). We previously showed that acetylation of H3K64 and H3K122 in the H3 core marks a subset of active enhancers in mESCs (Pradeepa et al., 2016). Whereas H3K115ac marks both active and repressed CpG island promoters, we find that H3K115ac is dynamically associated with active enhancers and is enriched within the nucleosome-depleted binding sites of key transcription factors. Its tight associated with chromatin accessibility suggests that H3K115ac could enhance detection of active enhancers when used in combination with other PTMs.

A distinct feature of H3K115ac is its association with fragile nucleosome. We show that H3K115ac is enriched in the NDRs associated with promoters, enhancers and CTCF sites. DNA associated with H3K115ac nucleosomes has higher A/T content than that associated with H3K27ac consistent with the enhanced MNase sensitivity of nucleosomes associated with A/T-rich sequences (Chereji et al., 2019). We confirm the association of H3K115ac with fragile nucleosomes by ChIP-seq of chromatin fractionated on sucrose gradients. In contrast, H3K27ac, H3K64ac and H3K122ac are depleted from fragile nucleosomes and NDRs.

Interestingly, H3K122 and H3K115ac exhibit distinct profiles despite their proximity within the structure of the nucleosome core suggesting specific and distinct mechanisms of deposition of these marks.

CTCF sites show an H3K115ac pattern distinct from promoters and enhancers as mononucleosomal H3K115ac signal is limited to +1 and −1 positioned nucleosomes flanking CTCF sites while the CTCF NDR is occupied by subnucleosomes marked with H3K115ac most likely in complex with CTCF protein. We note that weakly bound CTCF sites show a weaker nucleosome positioning but only in 3’ of the CTCF NDR and independent of TAD boundaries. Interestingly, CTCF N-terminus which interacts with cohesin, impeding or reversing cohesin-mediated loop extrusion (Li et al., 2020; Nora et al., 2020), aligns with 3’-end of the CTCF motif (Yang et al., 2023). Mechanisms of CTCF binding, orientation, H3K115ac and nucleosome remodeling may be interlinked.

Though nucleosome fragility has been associated with H3.3 and H2AZ histone variants (Jin et al., 2009), we show that H3K115ac is not restricted to H3.3. Our findings support the possibility that acetylation of H3K115 at the dyad axis of the nucleosome enhances the unwrapping or breathing of DNA from the core particle, or the release of histone subunits (Chatterjee et al., 2015, Zhou et al., 2019). The cBAF chromatin remodelling complex has recently been shown to act on fragile nucleosomes at mESCs enhancers to generate hemisomes that facilitate further Oct4 binding (Nocente et al., 2024). Given the impact of H3K115 acetylation on nucleosome stability in vitro (Manohar et al., 2009; Simon et al., 2011) and nucleosome disassembly by chromatin remodelers (Chatterjee at al., 2015), it is plausible that H3K115 acetylation facilitates the action of cBAF in further destabilising fragile nucleosomes at enhancers contributing to a dynamic chromatin state. Consistent with our observation of H3K115ac enrichment at repressed CGI promoters in mESCs, polycomb repressed promoters have also been shown to contain fragile nucleosomes attributed to BAF remodelling (Brahma & Henikoff. 2024). To the best of our knowledge, H3K115ac is the first histone modification enriched on a fragile nucleosome at the TSS of CGIs, at polycomb repressed promoters, at the TF binding sites of active enhancers, and at CTCF sites.

H3K115ac is therefore focussed at the heart of regulatory activity in the mammalian genome. In the future, it will be interesting to establish the functional significance of H3K115ac in modulating nucleosome structure, gene regulation and genome organization and to ascertain the value of H3K115ac profiling for the functional annotation of genomes.

## Materials and Methods

### Cell culture and neural differentiation

E14 mESC and its derived cells were cultured in GMEM-BHK21 medium (Gibco, #21710-082) supplemented with 10% heat inactivated foetal bovine serum (GIBCO, #A5256801 ), Penicillin-Streptomycin (GIBCO #15140-122), non-essential amino acids (GIBCO, #11140-035 ), sodium pyruvate (GIBCO, #11360-070), 2-Mercaptoethanol (GIBCO, #21985-023), GlutaMAX (GIBCO,#35050-087) and Leukaemia Inhibitory factor (in-house, Helin Lab) on 0.2% gelatine coated dishes or plates. Undifferentiated 46C Sox1-GFP mESCs (Ying et al., 2003) were cultured in GMEM-BHK21 medium (Gibco, #21710-025) supplemented with Leukaemia Inhibitory factor (103 units/ml. in- house), 10% foetal calf serum (Hyclone, #SH3007003), non-essential amino acids (Sigma, #M7145), sodium pyruvate (Sigma, #S8636), β-Mercaptoethanol (GIBCO, #31350010). Cell lines were validated and tested for mycoplasma at IGC, University of Edinburgh. 46C mESCs were differentiated to neural progenitor cells (NPCs) as described previously (Ying et al., 2003; Benabdallah et al., 2016). Cells were seeded at 3×10^6^ cells per 0.1% gelatin-coated T75 Corning flask 24 hours (h) prior to differentiation. Differentiation medium ((1:1DMEM/F12: Neuro basal medium (GIBCO, #31330-032 and 21103-049 respectively) supplemented with 0.5x B27 (Invitrogen, #17504044), 0.5x N2 (Invitrogen, #17502048), L-glutamine and 50mM 2-mercaptoethanol (GIBCO, #31350010)) was prepared fresh on the day of differentiation. On D0, ESCs were harvested, washed twice with PBS, twice with differentiation media, then seeded at a density of 1×10^6^ ESCs on 0.1% gelatin-coated T75 Corning flasks. Cells were cultured for 7 days with daily media changes after day 2. NPC differentiation was monitored visually through expression of the Sox1-GFP reporter.

### Generation of H3.3 KO mESC line

H3.3KO cells were generated by CRISPR/Cas9 targeting of DPY30-miniAID mESCs (Wang et al., 2023), using a combination of two sgRNAs to excise the exon 2 of H3f3a. mESCs were co-transfected with two eSpCas9 vectors (U6-sgRNA-eSpCas9(1.1)-T2A-mCherry and U6-sgRNA-eSpCas9(1.1)-T2A-GFP) encoding the sgRNA pair using Lipofectamine 3000 according to manufacturer’s instructions. GFP+/mCherry+ cells were single cell sorted 48h after transfection using a SONY MA900 cell sorter. Resulting clonal cell lines were screened by PCR genotyping for homozygous deletion and confirmed by immunoblotting. Clonal H3f3a knock out cells were used for the next round of H3f3b knockout use the same strategy to excise the exons 3 and 4 of H3f3b. Sequences of guide RNAs are in Supplementary table 5.

### Peptide Dot Blots

Custom peptides (ThermoFisher Scientific) with the sequence N-CAIHAK115RVTIMPK-C were synthesized as unmodified (H3K115), acetylated K115 (H3K115ac) and lysine to arginine mutation (H3K115R). For H3K122ac the peptide sequence was N-CGGVTIMPK122DIQLA-C. 2.5, 5,10 and 20 ng of peptide were spotted on a nitrocellulose membrane and allowed to dry at room temperature (r.t.). The blot was incubated in blocking buffer (5% non-fat dry milk in TBST: 20mM Tris, 150mM NaCl, + 0.15% Tween-20) for 30 minutes (mins), followed by primary antibody (1:1500 dilution of anti-H3K115ac, catalogue # PTM170, PTM Bio) in blocking buffer at r.t. before washing 3 x with TBST for 5 min each.

The blot was then incubated with a 1:10000 dilution of secondary antibody (HRP-linked anti-Rabbit IgG; Cell Signalling Technology, catalogue # 7074S) in blocking buffer for 45 mins at r.t. followed by washing as in previous steps. The blot was developed with HRP substrate (# 34096, ThermoFisher) and imaged on an ImageQuant LAS-4000 Imager. For peptide sequences see Supplementary table 6.

### Immunoblotting

Cells were washed with Dulbeccos-PBS, then lysed with 1x Laemmli buffer without 2-mercaptoethanol and Bromophenol blue. The lysates were boiled for 10min, the protein concentration quantified by BCA assay then adjusted, then 2-mercaptoethanol and Bromophenol blue was added. Lysates were boiled for 5min prior to loading. Proteins were run on a 4-20% TGX Gel (BioRad) and then transferred to a nitrocellulose membrane (LICOR). Total protein was visualised by Ponceau staining, then the membrane was blocked with 5% skimmed-milk in TBST, rinsed with TBST, and incubated with primary antibodies overnight, and washed with TBST, then incubated with secondary antibodies (LICOR), washed with TBST, and imaged on LICOR Odyssey M imager. For antibodies see Supplementary table 7.

### Native ChIP-seq with partial MNase digestion

mESCs and NPCs were harvested with Accutase (Lifetech) and washed twice with cold PBS. 5 million cells were permeabilized by resuspending in 100µL cold NBA buffer (5.5% Sucrose, 85 mM NaCl, 10 mM TrisHCl pH 7.5, 0.2 mM EDTA, 0.2 mM PMSF, 1 mM DTT, 1X Protease Inhibitors) and 100µL of NBB buffer (NBA buffer + 0.1% NP40) and incubated for 10 mins on ice. Nuclei were then collected by centrifugation at 1000g for 5 min at 4°C followed by washing with 200 µl NBR buffer (5.5% Sucrose, 85 mM NaCl, 10mM TrisHCl pH7.5, 3 mM MgCl2, 1.5 mM CaCl2, 0.2 mM PMSF, 1 mM DTT). Nuclei were resuspended in 100µl NBR buffer and incubated with 1µl of RNase-A (10mg/ml) for 5 min at r.t. followed by treatment with 4U MNase (Sigma) for 10 min at 20°C while mixing at 500 rpm. MNase was stopped by addition of an equal volume of ice cold 2x STOP buffer (215 mM NaCl, 5.5 % sucrose, 10 mM TrisHCl pH 8, 20 mM EDTA, 2 % Triton X100, 0.2 mM PMSF, 1 mM DTT, 2X Protease Inhibitors) and placing on ice. The reaction was diluted to 500µl total volume by adding NBR:STOP buffer and incubated on ice for 14 h to release digested fragments. Samples were centrifuged at 10000g for 10 mins at 4°C and solubilized chromatin was recovered as the supernatant. At this point, 10% of the chromatin was saved as Input. Dynabeads Protein-A (Invitrogen) were blocked and pH-calibrated as per the manufacturer’s protocol and 5 µl bed volume of beads were loaded with 2 µg of ChIP antibody (H3K27ac; Abcam-AB4729, H3K115ac; PTM BIO-PTM-170). Beads were added to the solubilized chromatin, supplemented with BSA to 0.1mg/ml and rotated for 4 h at 4°C. Beads were collected by a pulse spin and concentrated on a magnetic rack, washed 3x for 5 min each at r.t with 500 µL ChIP WASH buffer (10mM Tris pH 8, 2mM EDTA, 150mM NaCl, 1% NP-40, 1 % Na-Deoxycholate), followed by a single wash with TE buffer. Bead bound chromatin was eluted with 100µl elution Buffer (0.1 M NaHCO3, 1 % SDS) at 37°C while mixing at 1000 rpm for 15 mins. Input chromatin was diluted to 100 µl with elution buffer and the pH of Input and IP samples was adjusted by adding 7 µl of 2M TrisCl pH 6.8 and both were treated with 20 µg Proteinase-K for 2 h at 55°C while mixing at 1000 rpm. DNA was purified with Qiagen MinElute columns and sequencing libraries constructed using the NEBNext® Ultra™ II DNA library kit per the manufacturer’s instructions. Samples were sequenced for 50 cycles in paired- end mode on an Illumina HiSeq-2000.

### H3K115ac ChIP-qPCR with SNAP-ChIP® K-AcylStat panel as spike-in

SNAP-ChIP® K-AcylStat Panel from Epicypher (catalog # 19-3001) provides an equimolar pool of semi-synthetic nucleosome species each with a single lysine acetylation in combination with a unique DNA barcode. We used this panel as spike-in as per the manufacturer’s instructions and performed native MNase ChIP as described above. After purification of ChIP and Input DNA, qPCR was performed using primers complementary to the barcodes in the K-acyl panel as well as primer sets targeting the promoter and gene body of murine *Klf4* (Supplementary table 8) on a BioRad qPCR machine with 2x SYBR qPCR mix as per the manufacturer’s instructions. Recovery of different barcodes was computed as percent of total input chromatin (Lin et al., 2012).

### Sucrose gradient sedimentation of MNase digested chromatin

Chromatin from unfixed E14 mESCs as prepared as previously described (Nocente et al. 2024). Briefly, cells were permeabilized in 15 mM Tris-HCl pH 7.5, 15 mM NaCl, 5 mM MgCl2, 0.1 mM EGTA, 60 mM KCl, 0.3 M sucrose, 0.2% IGEPAL CA-630 and protease inhibitors for 10 min on ice and then centrifugated through an 8 ml sucrose cushion (15 mM Tris-HCl pH 7.5, 15 mM NaCl, 5 mM MgCl2, 0.1 mM EGTA, 60 mM KCl and 1.2 M sucrose) at 10,000g for 30 min. Nuclei were resuspended in MNase buffer (20 mM Tris-HCl pH 7.5, 20 mM NaCl, 2 mM CaCl_2_, 4 mM MgCl_2_, and 15 mM KCl), and digested for 10 min at 37 °C with 1.4 Kunitz units of MNase (New England Biolabs, 200 Kunitz units per μl) per 1 x 10^6^ cells. MNase was stopped by addition of EDTA to 10 mM and EGTA to 20 mM. After incubation for 1 h on a rotative agitator at 4 °C, chromatin was released by passing the suspension 13 times through a 26-gauge needle, and insoluble material removed by centrifugation. Solubilized chromatin was separated into 500 μl batches derived from approximately 8 x 10^7^ cells. Each batch was loaded onto a 9.9 ml 10-30 % sucrose gradient containing 10 mM Tris-HCl pH 7.5, 10 mM EDTA, 20 mM EGTA, and 80 mM NaCl centrifugated for 16 h at 197,000g, 4 °C, using a Beckman Coulter SW 41 swinging-bucket rotor. 20 fractions of 500 μl were collected from the top to the bottom of the tube.

As previously described (Nocente et al., 2024), fractions 5 to 8 contain subnucleosomal particles of increasing size, fractions 9-10 are enriched in fragile nucleosomes, and fractions 11 to 13 are composed of stable mononucleosomes. Fractions corresponding to each of these species were pooled, the chromatin quantified by Qbit dsDNA HS assay and equal amounts of chromatin were taken from each pool and topped up to 1 ml using low salt buffer (10 mM Tris-HCl pH 7.5, 10 mM EDTA, 20 mM EGTA, and 80 mM NaCl) to ensure identical chromatin concentrations in all ChIP reactions. Equal amounts of Spike-In chromatin (from MCF7 cells) were added in the ratio of 1:30, and input samples (10% of total) were saved.

### ATAC-seq sample preparation

ATAC-seq libraries were prepared from 5×104 cells as previously described (Buenrostro et al., 2013) with some modifications (Wong et al 2023). We used 15 mins incubation on ice in the nuclei preparation step and the Tn5 reaction was performed in 50 µl of custom transposition buffer (10 mM Tris pH 8, 5 mM MgCl2 and 10% dimethylformamide) with 2.5 µl Tn5 transposase (Illumina, 20034197) at 37°C while mixing at 1000 rpm for 30 mins. The reaction was stopped and DNA purified with a MinElute PCR Purification Kit (Qiagen, 28004). To minimize PCR duplication, we used 20% of tagmented DNA for a test qPCR reaction to determine the minimum number of PCR cycles required to amplify the DNA without entering the plateau phase. The rest of the tagmented DNA was then amplified in identical conditions of dilution and PCR cycles. PCR reactions were purified and unused adaptors removed with an equal volume of AMPure XP beads (Beckman, A63880) following the manufacturer’s protocol and eluted in 20 µl Tris pH 7.8. Libraries were quantified using a Qubit assay (Invitrogen, Q32851) and were selected for the presence of a clear nucleosomal ladder pattern on a Bioanalyzer High Sensitivity assay (Agilent). Samples were sequenced for 50 cycles in paired-end mode on an Illumina HiSeq-2000.

### 4SU sequencing (4SU-seq)

4SU-seq was performed as described previously (Boyle et al., 2020). 4-thiouridine (4SU; Sigma T4509) was added to cell cultures to a final concentration of 500 µM and incubated for 20 mins at 37°C. Cells were collected by trypsinization, and total RNA purified from 5×10^6^ cells with Trizol (Invitrogen, 15596026) and DNase-treated with a Turbo DNA-free kit (Invitrogen, AM1907M). Thirty micrograms of total RNA were biotinylated using 60 µg of Biotin-HPDP (Pierce, 21341, 1mg/ml stock in dimethylformamide) in a 300 µl reaction in biotinylation buffer (10 mM TrisHCl pH 7.4, 1 mM EDTA) for 90 mins at r.t. Uncoupled biotin was removed by chloroform extraction (1:1 volume) and labelled RNA precipitated using isopropanol. RNA was resuspended in 100 µl RNAse-free water and incubated with an equal volume of µMacs streptavidin beads (Miltenyi 130-074-101) for 15 mins at r.t. Beads were captured on a pre-calibrated µMacs column, followed by 3 washes with 900 µl of Wash Buffer pre-heated to 65°C followed by 3 washes at r.t. RNA was eluted with 100 mM DTT and purified using an RNeasy MinElute cleanup kit (Qiagen 74204) as per the the manufacturer’s instructions. Ribosomal RNA was removed using a RiboMinus eukaryote system v2 kit (Ambion, A15027) and stranded libraries constructed using the NEBNext Ultra II directional RNA library preparation kit (NEB, E7760) following the protocol for ribosome-depleted RNA and with an 11-min RNA fragmentation step according to the manufacturer’s instructions. Libraries were amplified, purified and sequenced in paired-end mode as described for ATAC-seq.

### Data Analysis

#### Quality assessment and pre-processing of sequencing data

Publicly available datasets were downloaded from the NCBI SRA repository with SRA toolkit (fastq-dump). We used FASTQC v0.11.7 for quality control metrics. Filtering of poor-quality reads and trimming of adaptor sequences was performed using fastp (Chen et. al., 2018) using default settings.

#### ChIP-seq analysis

***Single-end ChIP libraries:*** Filtered and trimmed reads were aligned to the mm10 assembly of the mouse genome using bowtie (Langmead and Salzberg, 2012) and uniquely mapped reads were kept. Alignments were converted to BAM format and sorted using SAMtools (Li et al., 2009) and PCR duplicates removed using Picard (https://broadinstitute.github.io/picard/). Bedtools (Quinlan and Hall, 2010) was used to create BED files followed by conversion into HOMER (Heinz et al., 2010) tag directory format (makeTagDirectory) using default parameters. We called peaks using HOMERs findPeaks command with options -F 2 -localSize 50000 -size 150 -minDist 300 -fdr 0.01 – region. UCSC compatible bigwig files normalized to sequencing depth were generated using makeUCSCfile with parameters: -bigWig -style chipseq – fragLength 150.

***Paired-end ChIP libraries:*** Filtered and trimmed reads were aligned to the mm10 genome using bowtie2 with options (--no-discordant --no-mixed --no-unal -X 2000) followed by conversion to BAM format and removal of PCR duplicates as above. Read pairs were converted into ChIP fragments using bedtools bamtobed –bedpe and fractionated into sub-nucleosomes (<150 bp) and mono-nucleosomes (150-225 bp). Fragment coverage was computed at base-pair resolution with bedtools genomeCoverageBed and coverage normalized to sequencing depth and scaled to 107. Coverage data was converted to bigwig format using wigTobigWig (Kent utilities, UCSC). We computed Pearsons correlation between two replicates in 10Kb windows across the genome using the normalized coverage data and replicates were merged based on high correlation (Pearsons coeff. > 0.9).

#### Differential analysis of ChIP-seq data

To compute significant changes in H3K115ac during ESC to NPC differentiation, we counted reads within 500 bp upstream and downstream of TSS and used these for differential binding in DESeq2 (Love et. al., 2014). Differential binding is defined as fold change >=2 and pAdj <=0.01.

#### Spike-in normalization of ChIP-seq libraries

A hybrid genome was created by catenating mouse (mm10) and human (hg38) genome sequences with species-specific chromosome names and a genome index was prepared using *bowtie2-build* command from bowtie2 with default parameters. Reads were preprocessed and aligned to this hybrid genome index as described above for paired-end ChIP libraries. Reads aligning to mouse and human genome were separated and counted for each sample using samtools. A scaling factor was computed for each ChIP and input library as described earlier (Bressan et al., 2024):

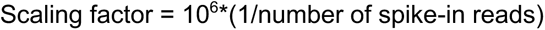

Read coverage was computed at 10 bp resolution across the mouse genome and multiplied by the scaling factors derived above.

#### ATAC-seq analysis

Trimming, filtering, and alignment steps are as described for paired-end ChIP-seq above. The 5′ ends of the reads were offset by +4 bases for reads on the Watson strand, and by −5 bases for the reads on the Crick strand, to reflect the exact location of Tn5 insertion. Peaks were called using *macs2 callpeak–nomodel–extsize 150–shift −75 -g ‘mm’ -p 0.01*.

Irreproducible discovery rate (IDR, ≤0.05, Li et al., 2011) was used to select reproducible peaks from two biological replicates. For visualization, a 20 bp fragment was centered on each Tn5 insertion site and its coverage was calculated at single-base resolution in the whole genome. Scaling and normalization steps were as for paired-end ChIP-seq above.

### 4SU-seq analysis

Trimming, filtering and alignment steps are as described for paired-end ChIP-seq. Alignments were converted into HOMER tag directory format with options suitable for stranded libraries *-format sam -flip –sspe*. Strand-specific genome wide coverage (bigwig format) was computed with parameters: *makeUCSCfile –bigwig -fsize 1e20 -strand + (or −) - norm 1e8.* Read coverage for promoters was computed using HOMER’s *annotatePeaks.pl* with parameters: *-size “given” -noann -nogene -len 0 -strand both -norm 0.* Read counts were then converted into reads per kilobase per million mapped reads (RPKM). Differential expression analysis was carried out at gene level using limma package (Ritchie et. Al., 2015).

#### MNase-seq data processing and nucleosome calls

Classical MNase-seq (fragments within mononucleosomal range) data for mESCs was downloaded from NCBI SRA (GSE59062, West et. al., 2014). Trimming, filtering, and alignment steps are as described for paired-end ChIP-seq. High quality alignments (MAPQ >=30) from two biological replicates were merged using *SAMtools*. Nucleosome peaks were called using the DANPOS3 algorithm (Chen et. al., 2015) with default parameters. Single base resolution wig files from DANPOS3 were converted to bigwig files for visualization in the UCSC genome browser and plotting coverage using *deeptools* (Ramírez et. al., 2016).

#### Promoter, CGI and TSS definitions

Gene annotations were downloaded from the UCSC table browser for genome build: mm10, track: NCBI RefSeq, table: UCSC RefSeq (refGene). For a given TSS (coding and non-coding), 1000 bp upstream and downstream were added and used as a promoter region to filter out promoter proximal peaks in downstream analyses. To create a reference set of unique protein-coding TSSs without duplicated entries, we collapsed the transcripts with identical TSS location. For use in V-plots, a more conservative set of TSSs was defined such that: i) for genes with multiple TSSs, a single TSS with highest 4SU-seq count in mESCs was retained, ii) genes with single TSSs were included, iii) TSSs were filtered out if there was another coding/non-coding TSS within 1 Kbp. This ensured that the local and directional observations in our analyses are not affected by transcriptional activity in the vicinity. Association of promoters with CGIs was defined based on a CGI located within 500bp upstream or downstream of the TSS.

#### Random forest model

ATAC-seq accessible non-promoters were overlapped with H3K115ac, H3K122ac, and H3K27ac peaks and classified as H3K115ac-positive or negative. Peaks from the cistromeDB (Mei et al., 2017) and ReMap2022 (Hammal et al., 2022) were filtered as described previously (Friman et al., 2023). The PeakPredict package (https://github.com/efriman/PeakPredict) command *overlap_peaks* was run with settings *‘--predict_column H3K115ac --model RandomForestClassifier --balance --shap’,* where balancing downsamples the groups to have the same size prior to splitting into test and training sets. The PeakPredict package implements scikit-learn (Pedregosa et al., 2011) and SHAP values (Lundberg and Lee, 2017).

#### CTCF ChIP-seq peaks and motif calls

Analysis of nucleosome positioning around CTCF sites using CTCF ChIP-seq peaks as reference point suffers from low resolution. To mitigate this, we intersected CTCF peaks (Mas et al., 2018) in the mESC genome with CTCF motif calls and selected peaks with a single CTCF motif. We used these motif coordinates as reference point to plot histone acetylation data. Replicate merged CTCF ChIP-seq peaks for mESCs were downloaded from Remap2022 (Mas et al.,2018; Hammal et. al., 2022; GSE99530). ChIP-seq peak scores were used to split data into quartiles in R. The mouse genome was scanned for concensus CTCF motifs (JASPER: MA0139.1) using *matchPWM* in the Biostrings package in Bioconductor. We removed any peaks that overlapped with > one CTCF motif and CTCF binding scores divided into quartiles.

#### Contour plots

For TSS and CTCF motifs we plotted signed distance (x-axis) and fragment length (y-axis). We defined equal number of high-density regions as contours using the package ggdensity in ggplot2 (Otto and Kahle, 2023). Contours refer to different levels of fragment density as shown by color density. Each contour is associated with a probability (%) with which it is bound to data.

#### Datasets

The following publicly accessible datasets were used in this study; NCBI GEO GSE66023 (mESC H3K122ac, H3K27ac), GSM1003756 (mESC H3K4me3), GSM1003750 (mESC H3K4me1), GSM1276707 (mESC H3K27me3), GSE59062 (mESC MNase), GSE115774 (mESC 4SU-seq) and GSE78910 (mESC H3.3 turnover). Sequencing data generated in this study are submitted to NCBI GEO with the Accession number GSE246191.

## Author Contributions

DS, YK, MS, ZF and HW: Designed and performed the experiments; YK, EF, RI: Performed data analyses; DS, WB, YK: conceived the project; WB , K H , M G re s o u rc e d and ma n a g e d t h e p r o j e c t ; YK, WB: wrote the manuscript; all authors edited the manuscript.

## Acknowledgements.

We thank Nick Gilbert for discussions. This work has made use of the resources provided by the Edinburgh Compute and Data Facility (ECDF) (http://www.ecdf.ed.ac.uk/) for sequencing data analysis and the Wellcome Trust Clinical Research Facility, Edinburgh (https://clinical-research-facility.ed.ac.uk/) for sequencing.

## Funding statement

DS was supported by a Newton Fellowship from the Royal Society. Work in the WAB lab is funded by MRC University Unit grant MC_UU_00035/7. E.T.F. was supported by the Swiss National Science Foundation (P500PB_206805).

---

**Figure S1.**
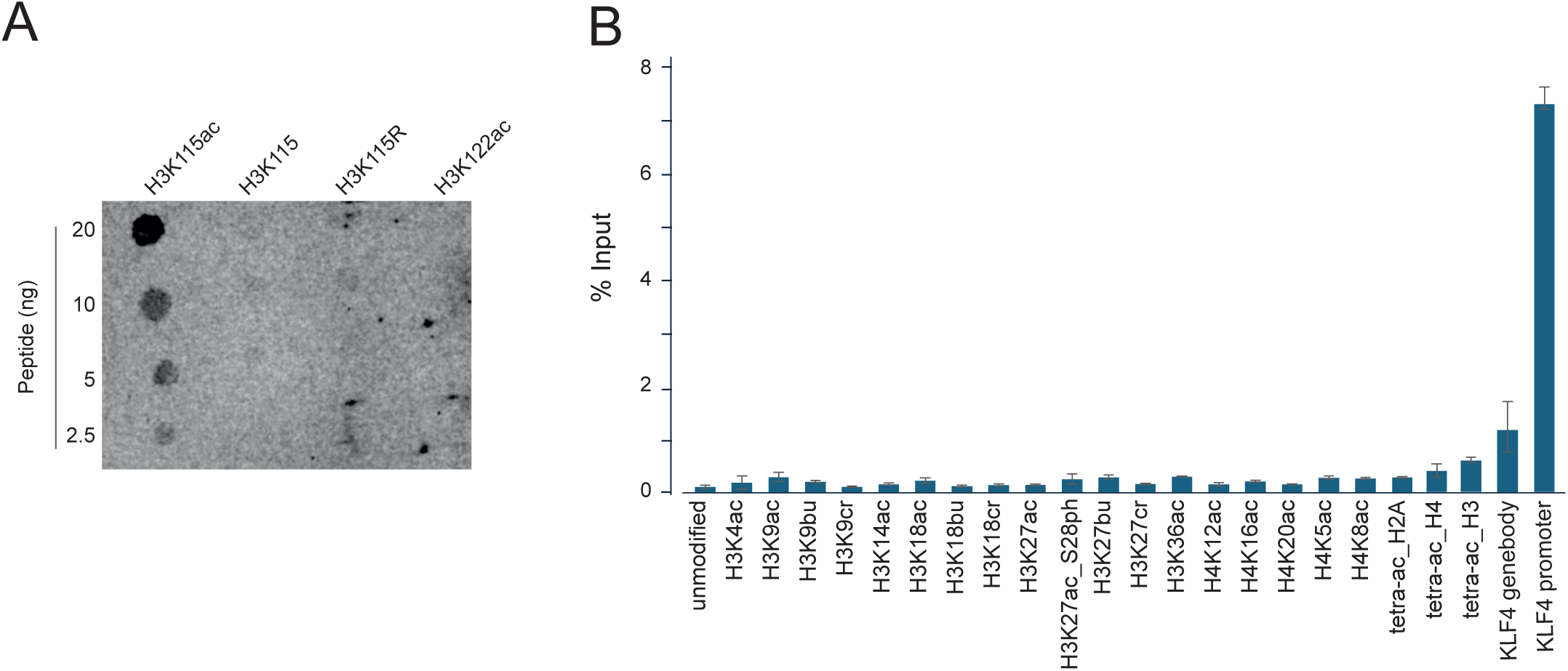
H3K115ac antibody specificity. Related to Figure 1. A) Dot blot for specificity of H3K115ac antibody against unmodified H3K115 and H3K115R peptides and acetylated H3K115 and H3K122 peptides. B) ChIP-qPCR with H3K115ac antibody from mESC chromatin spiked with equimolar amounts of bar-coded nucleosome species modified as indicated; acetyl (ac), butyryl (bu), crotonyl (cr), phopsho (ph) from the SNAP-ChIP K-acyl-stat panel. The gene body and promoter of *Klf4* are included as endogenous targets. Error bars indicate standard deviation from three technical replicates for each of two biological replicates.

**Figure S2.**
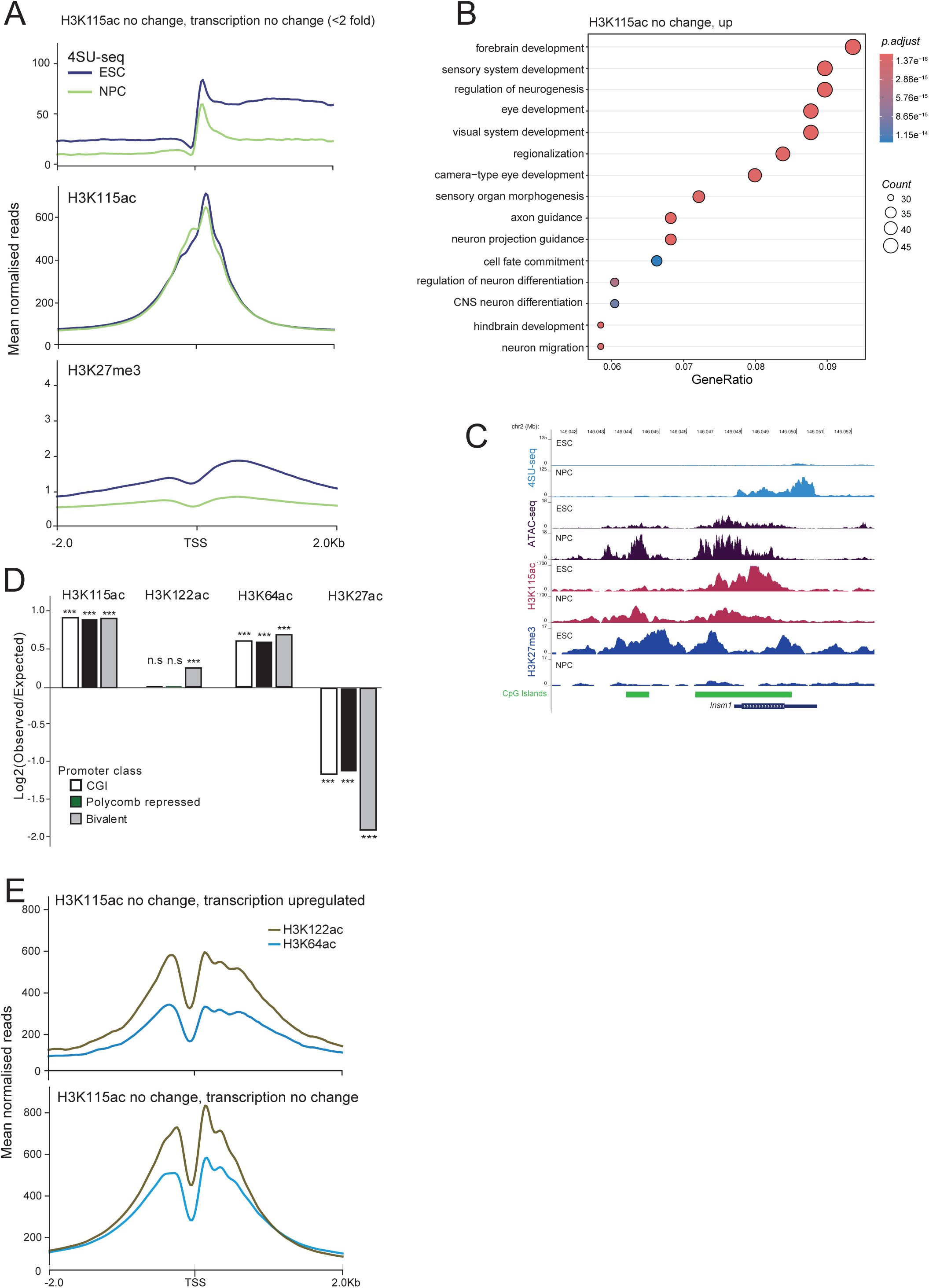
H3K115ac dynamics during differentiation. Related to Figure 2. A) Aggregate profile plots for 4SU-seq, and ChIP-seq data for H3K115ac and H3K27me3 (Mikkelsen et al., 2007) at promoters with no significant change in H3K115ac occupancy, or transcription (<2 fold) during differentiation to NPCs. B) Gene ontology analysis for the background set of genes in Figure 2C. Top 15 biological processes are shown. C) UCSC genome browser screenshot showing 4SU-seq, ATAC-seq, and ChIP-seq data for H3K115ac, and H3K27me3 (Mikkelsen et al., 2007) in mESCs and differentiated NPCs at the *Insm1* locus. CpG islands (CGI) are indicated. Genome co-ordinates (Mb) are from the mm10 assembly of the mouse genome. D) Enrichment (Log2 observed/expected) of H3K115ac, H3K122ac, H3K64ac and H3K27ac ChIP-signal with polycomb target promoters (H3K27me3) in mESCs. *** p value (Fishers) <0.01. n.s. not significant (p>0.05). E) H3K64ac and H3K122ac mESC ChIP-seq profiles at the gene sets from Figure 2C and Figure 2SA that show no change in H3K115ac ChIP signal during ESC to NPC differentiation and that are either transcriptionally up-regulated (Figure 2C), or that show no change in transcription upon differentiation (Figure 2SA) (ChiP-seq data from Pradeepa et. al., 2016).

**Figure S3.**
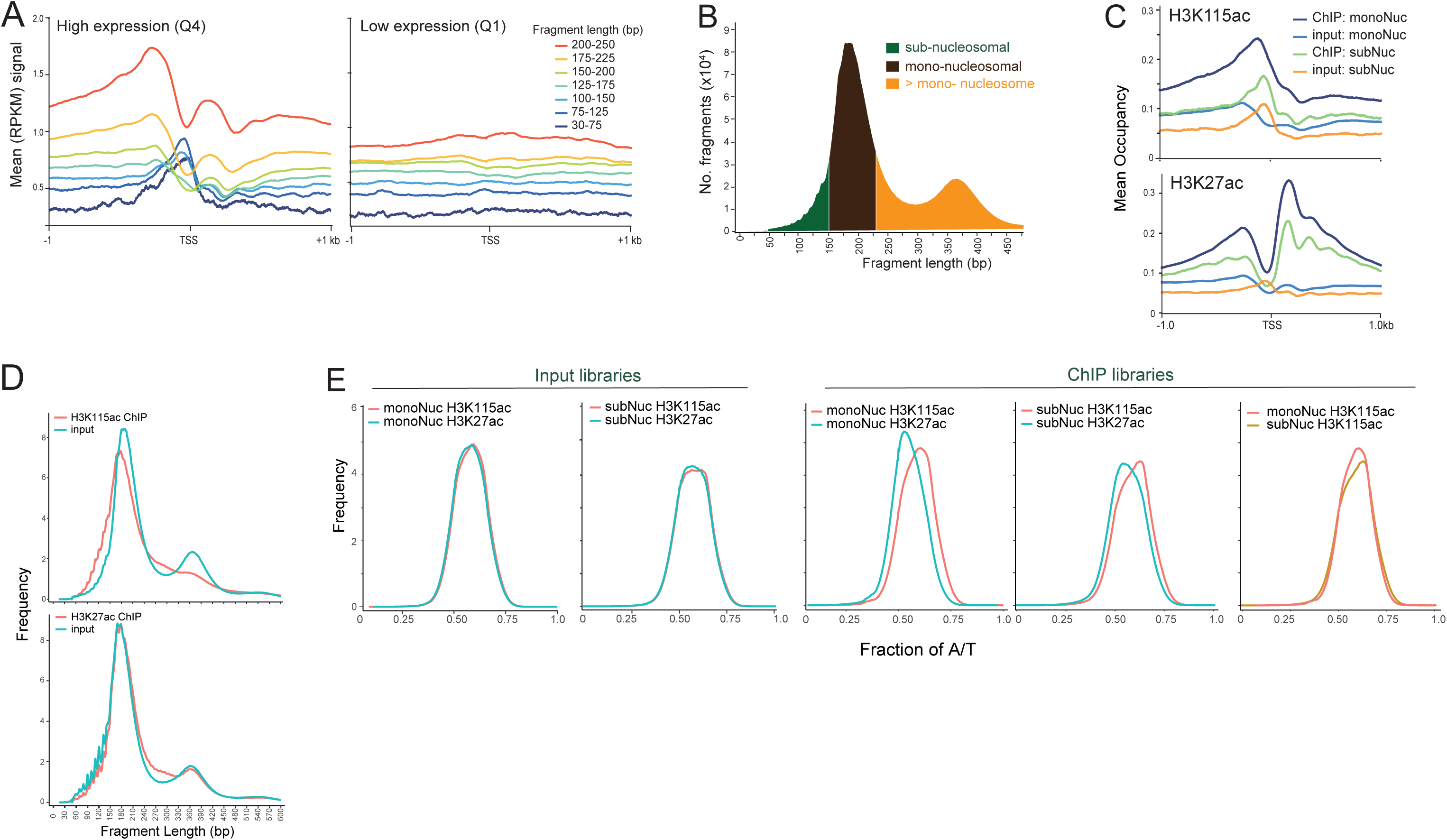
H3K115ac is associated with sub-nucleosome sized fragments. Related to Figure 3. A) Mean coverage of fragments in MNase-digested input libraries for the most highly active TSSs (4SU Q4) and minimally active TSSs (Q1) in mESCs binned into different fragment lengths (bp). B) Schematic of selected fragment lengths to define sub-nucleosomes and mono-nucleosomes from input libraries. C) Mean occupancy of H3K115ac (top) or H3K27ac (below) ChlP-seq from mESCs plotted around (+/-1kb) TSS. Sub-nucleosomal and mono-nucleosomal signals are plotted with together with the profile from their respective input libraries. D) Distribution of fragment lengths (bp) from paired-end libraries for; (top) native MNase ChIP-seq of H3K115ac and its input sample, (bottom) H3K27ac ChIP-seq libraries. Distribution is scaled to library size and the Wilcox test was used to calculate significance (Supplementary table 3). E) Density profiles of A/T content for Input and H3K115ac and H3K27ac ChIP-seq of libraries showing mono-nucleosomes (monoNuc) and (right) sub-nucleosomes (subNuc). A/T content does not differ between different input libraries (Wilcox test, p > 0.01), but H3K115ac marked subnucleosomes have a higher A/T content than subnucleosomes marked with H3K27ac (Wilcox test, p <0.01), H3K115ac marked mono-nucleosomes have higher A/T content than those with H3K27ac (Wilcox test, p <0.01). The A/T content of sub- and mono-nucleosomal particles marked with H3K115ac are not significantly differentt (Wilcox test, p > 0.01). Statistical data in Supplementary table 3.

**Figure S4.**
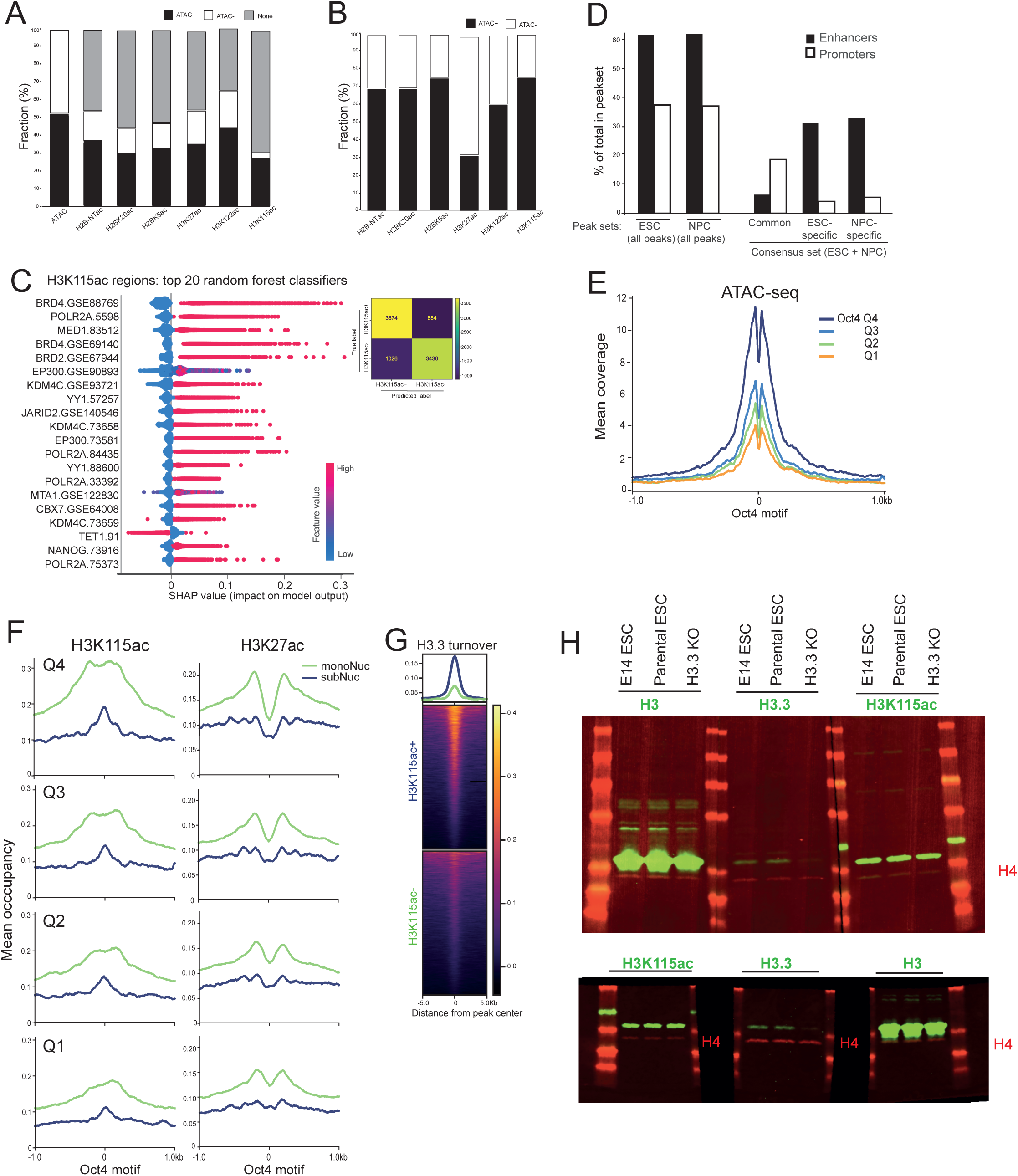
H3K115ac correlates with regulatory activity at mESC enhancers. Related to Figure 4. A) Overlap of STARR-seq +ve sequences (Peng et al., 2020) with sites corresponding to; *open* (ATAC+) or *closed* (ATAC-) chromatin sites in the mESC genome, and for different mESCs histone aceyl-lysine peaks. ‘None’ indicates STARR-seq active sequences with no overlap with the indicated post translational modification class. H2B-NTac refers to regions marked with at least one of the acetyl-lysines in the N-terminal of histone H2B (K5, K11, K12, K16 and K20, Narita et al., 2023). B) the proportion of peaks for different histone acetylation marks that are defined as open or closed by ATAC-seq in mESCs. C) A Random Forest model was trained to predict H3K115ac-positive versus -negative peaks (but marked by H3K27ac and/or H3K122ac) based on ChIP-seq overlap. SHAP values were calculated to determine the impact on the model (top 20 features shown, left). The prediction confusion matrix is shown for the test data (right). D) Bar plot showing % of promoter- or enhancer-associated peaks in mESC and NPCs. Nearly 33% of all H3K115ac peaks are associated with promoters in both ESCs and NPCs with 70% of them common between ESCs and NPCs. In contrast, the majority of dynamic H3K115ac peaks are associated with enhancers (ESC-specific; n=1881, NPC-specific; n=2811). E) ATAC-seq signal around Oct4-occupied enhancers in mESCs divided into quartiles (Q1-Q4, low to high) of Oct4 ChIP-seq peak strength. F) H3K115ac (left) and H3K27ac (right) paired-end ChIP signal around Oct4-occupied enhancers. ChlP-seq reads are separated according to fragment length into mononucleosomes (green; monoNuc) and subnucleosomes (blue; subNuc). G) Heatmap depicting H3.3 turnover index (Deaton et al., 2016; GSM2080325_TI_ES.wig in GEO accession GSE78910) centered for H3K115ac-+ve and -ve regions. H) Immunoblotting for H3, H3.3 and H3K115ac (green) and H4 (red) on whole cell extracts from E14 mESCs, H3.3 knock-out ESCs and the parental ESC line. Red size markers are also shown. Top panel is complete gel corresponding to Fig 4D, lower panel is a biological replicate.

**Figure S5.**
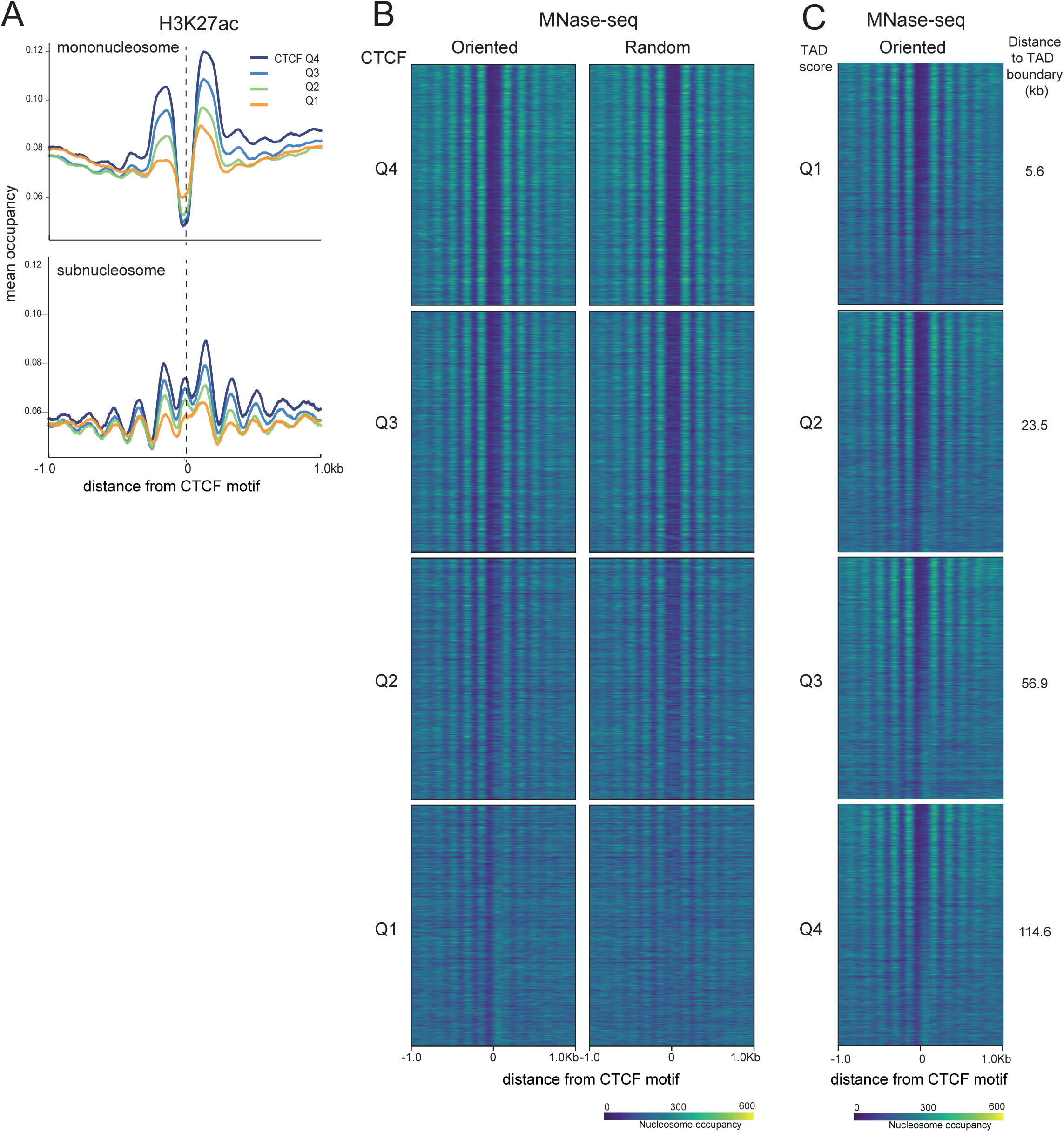
H3K27ac and H3K115ac relative to CTCF binding sites and motif orientation. Related to Figure 5. A) H3K27ac ChlP-seq signal from mononucleosomal (top) and subnucleosomal (bottom) sized fragments in mESCs around CTCF motifs across the four quartiles of CTCF ChlP-seq peak strength. All CTCF motifs are oriented from 5’ to 3’ (left to right). B) Heatmap of MNase-seq data ranked by quartiles of CTCF occupancy with CTCF motifs oriented in the same (left) direction or randomised orientation (middle). C) As in (B) but divided into quartiles (Q1-Q4) of distance to the closest TAD boundary in mESCs (TAD boundaries from Bonev et al., 2017). Median distance (kb) to TAD boundary for each quartile is shown on the right; minimum and maximum distances are shown in Supplementary table 4. Within each TAD quartile, sites are sorted by CTCF occupancy in descending order.

## Notes

### Competing Interest Statement

The authors have declared no competing interest.

### Summary of Updates

We now provide a biological replicate for the Western Blot (new FigS4H) together with an image of the whole gel for the data in Fig 4D. We have corrected some minor formatting issues and toned down a few claims in the text.

