## Supplementary tables for "Acetylation of H3K115 is detected associated with fragile nucleosomes at CpG island promoters and active regulatory sites"

**Supplementary Table 1**

| Replicate | Pearsons Correlation | Sequencing format |
|---|---|---|
| H3K27ac_mESC_peChIP | 0 9945 | Paired end |
| H3K27ac_mESC_peChIP_Input | 0 9992 | Paired end |
| H3K115ac_mESC_peChIP | 0 9905 | Paired end |
| H3K115ac_mESC_peChIP_Input | 0 9993 | Paired end |
| H3K115ac_mESC_seChIP | 0 9684 | Single end |
| H3K115ac_NPC-day7_seChIP | 0 904 | Single end |
| ATAC-seq_mESC | 0 9369 | Paired end |
| ATAC-seq_NPC-day7 | 0 9528 | Paired end |

**Supplementary Table 2**

**Statistical data for Figure 2B**

**Fisher's exact test (association of gain/loss of H3K115ac with transcriptional changes on promoters classified based on association with CpG islands)**

**All promoters**

| Categories | nuber of genes | log2 (observed / expected) | p-value | Interpretation |
|---|---|---|---|---|
| Gain ~ up | 517 | 1 00 | 2 02E-57 | Significant positive enrichment |
| Gain ~ down | 215 | -0 42 | 0 000102 | Significant negative enrichment |
| Loss ~ up | 163 | -0 15 | 0 769 | Negative enrichment, not significant |
| Loss ~ down | 524 | 1 39 | 2 93E-111 | Significant positive enrichment |

**Promoters with CGI**

| Categories | nuber of genes | log2 (observed / expected) | p-value | Interpretation |
|---|---|---|---|---|
| Gain ~ up | 374 | 0 78 | 1 21E-25 | Significant positive enrichment |
| Gain ~ down | 187 | -0 28 | 0 000305 | Significant negative enrichment |
| Loss ~ up | 150 | -0 06 | 0 337 | Negative enrichment, not significant |
| Loss ~ down | 385 | 1 24 | 4 44E-59 | Significant positive enrichment |

**Promoters without CGI**

| Categories | nuber of genes | log2 (observed / expected) | p-value | Interpretation |
|---|---|---|---|---|
| Gain ~ up | 168 | 1 80 | 2 33E-52 | Significant positive enrichment |
| Gain ~ down | 28 | -1 06 | 0 00000308 | Significant negative enrichment |
| Loss ~ up | 13 | -1 19 | 0 00053 | Significant negative enrichment |
| Loss ~ down | 139 | 1 95 | 1 93E-51 | Significant positive enrichment |

**Fisher's test for association with bivalent genes from Seneviratne et. al., 2024**

| Categories | log2 (observed / expected) | p-value | interpretation |
|---|---|---|---|
| ncK115_down | -0 13 | 4 4E-01 | ns |
| ncK115_nc | 0 14 | 5 7E-14 | Mild but significant enrichment |
| ncK115_up | 1 95 | 1 6E-91 | Strong and significant enrichment |
| upK115_nc | 0 89 | 4 5E-18 | Strong and significant enrichment |
| upK115_up | 1 95 | 6 2E-22 | Strong and significant enrichment |
| downK115_down | 0 16 | 5 8E-01 | ns |
| downK115_nc | 1 23 | 9 5E-49 | Strong and significant enrichment |

**Fisher's test for association with polycomb target promoters (related to Figure S2D)**

**All promoters with a CGI**

| PTM | log2 (observed / expected) | p-value | interpretation |
|---|---|---|---|
| H3K115ac | 0 911600342 | 0 | positive and significant enrichment |
| H3K122ac | 0 006175312 | 0 4399634 | mild but insignificant enrichment |
| H3K64ac | 0 602910054 | 0 | positive and significant enrichment |
| H3K27ac | -1 098831002 | 0 | negative and significant enrichment |

**Bivalent genes (Seneviratne et. al., 2024)**

| PTM | log2 (observed / expected) | p-value | interpretation |
|---|---|---|---|
| H3K115ac | 0 9200524 | 2 79E-280 | positive and significant enrichment |
| H3K122ac | 0 2556087 | 1 16E-46 | mild and significant enrichment |
| H3K64ac | 0 6928932 | 2 75E-222 | positive and significant enrichment |
| H3K27ac | -1 8731599 | 0 00E+00 | negative and significant enrichment |

**all genes bound with EZH2**

| PTM | log2 (observed / expected) | p-value | interpretation |
|---|---|---|---|
| H3K115ac | 0 925373461 | 0 | positive and significant enrichment |
| H3K122ac | 0 008865189 | 0 4047265 | mild but insignificant enrichment |
| H3K64ac | 0 612832156 | 0 | positive and significant enrichment |
| H3K27ac | -1 137062504 | 0 | negative and significant enrichment |

**Supplementary Table 3**

Statistical data for Figure S3D

**p values (Wilcox test)  fragment length distributions**

| Distribution_1 | Distribution 2 | p. Value | related figFigure 3E |
|---|---|---|---|
| H3K115ac ChIP Replicate 1 | H3K115ac Input Replicate 1 | 8 03E-132 | Figure S3D, top panel |
| H3K115ac ChIP Replicate 2 | H3K115ac Input Replicate 2 | 1 23E-67 | Figure S3D, top panel |
| H3K27ac ChIP Replicate 1 | H3K27ac Input Replicate 1 | 3 96E-08 | Figure S3D, bottom panel |
| H3K27ac ChIP Replicate 2 | H3K27ac Input Replicate 2 | 0 2946652 | Figure S3D, bottom panel |

Statistical data for Figure S3E

**p values (Wilcox test)  AT content distributions**

| Distribution_1 | Distribution 2 | p. Value | related figure |
|---|---|---|---|
| ChIP_subNuc_H3K115ac | ChIP_subNuc_H3K27ac | 1 97E-08 | Figure S3E |
| ChIP_monoNuc_H3K115ac | ChIP_monoNuc_H3K27ac | 3 49E-23 | Figure S3E |
| ChIP_monoNuc_H3K115ac | ChIP_subNuc_H3K115ac | 0 3901559 | Figure S3E |
| Input_monoNuc_H3K115ac | Input_monoNuc_H3K27ac | 0 1582149 | Figure S3E |
| Input_subNuc_H3K115ac | Input_subNuc_H3K27ac | 0 3641454 | Figure S3E |

**Supplementary Table 4**

**Quartiles of distances between CTCF motifs and the closest TAD boundary (Related to Figure S5C)**

| Quartile | Distance to the nearest TAD Boundary (base pairs) | | |
|---|---|---|---|
| | Minimum | Maximum | Median |
| Quartile_1 | 0 | 12 948 | 5596 |
| Quartile_2 | 12 949 | 38 164 | 23499 |
| Quartile_3 | 38 173 | 80 693 | 56852 |
| Quartile_4 | 80 711 | 2 620 841 | 114574 |

**Supplementary Table 5**

**sgRNAs used in generation of H3.3 KO line (Figure 4D)**

| | |
|---|---|
| H3f3a_sgRNA1_F | CACCgattaatttccagatttggg |
| H3f3a_sgRNA1_R | AAACcccaaatctggaaattaatc |
| H3f3a_sgRNA2_F | CACCgagaccagtaaagttcccaa |
| H3f3a_sgRNA2_R | AAACttgggaactttactggtctc |
| H3f3b_sgRNA1_F | CACCgaccgcggtcctaaaagcca |
| H3f3b_sgRNA1_R | AAACtggcttttaggaccgcggtc |
| H3f3b_sgRNA2_F | CACCgccattccagagattggtga |
| H3f3b_sgRNA2_R | AAACtcaccaatctctggaatggc |

**Genotyping primers used in generation of H3.3 KO line**

| Gene | Forward | Reverse |
|---|---|---|
| H3f3a | gtgtttgtggcttcgttcatt | agccctgccttctaaacatac |
| H3f3b | gaagagctccctagtgctaaac | cagtcagtcactcttcccattc |

**Supplementary Table 6**

**Peptide Sequences for dot blots (Figure S1A)**

| | |
|---|---|
| H3K115 | CAIHAKRVTIMPK |
| H3K115R | CAIHARRVTIMPK |
| H3K115ac | CAIHAK[Ac]RVTIMPK |
| H3K122ac | CGGVTIMPK[Ac]DIQLA |

**Supplementary Table 7**

**Antibodies used in immunoblotting**

| Target | Catalog No. | Manufacturer | Host | dilution |
|---|---|---|---|---|
| H3K115ac | PTM170 | PTM Bio | Rabbit | 1/1000 |
| Histone H3.3 | 09-838 | Millipore | Rabbit | 1/1000 |
| Histone H3 | AB1791 | Abcam | Rabbit | 1/5000 |
| Histone H4 | CST2935 | CST | Moouse | 1/3000 |
| IRDye® 800CW Goat anti-Rabbi | 926-32211 | LI-COR | Goat | 1/20,000 |
| IRDye® 680RD Goat anti-Mouse | 926-68070 | LI-COR | Goat | 1/10,000 |

**Supplementary Table 8**
**Primer Sequences (Figure S1B)**

| Primer ID | Sequence |
|---|---|
| Universa_Forward | CGTCGAACGCGCGATAT |
| Unmodified_Rev | TCGAATCGTTCGCGTAATC |
| H3K4ac_Rev | CGCGATTCGCGTAATACG |
| H3K9ac_Rev | ACGCGAAACGACGAATC |
| H3K9bu_Rev | GAATCGTCGACGCGTATA |
| H3K9cr_Rev | ATTCGCGCGTACGTATAC |
| H3K14ac_Rev | CGAAATTCGTATACGCGTCG |
| H3K18ac_Rev | CGCGTAACGACGTACC |
| H3K18bu_Rev | GCGACGCGTAATCGA |
| H3K18cr_Rev | GCGATCGCGCGAATA |
| H3K23ac_Rev | ACGCGTCGAAACGATTA |
| H3K27ac_Rev | CGTATACGCGCGACA |
| H3K27bu_Rev | TACGCGCGAATTTACGTC |
| H3K27cr_Rev | GAACGTTCGTCGACGAT |
| H3K36ac_Rev | TCGCGCGTACGAAAC |
| H4K5ac_Rev | CGAATTTCGCGCGTATTAC |
| H4K8ac_Rev | CGCGAACTATCGTCGATTC |
| H4K12ac_Rev | ATACGACGAGATAGTCGACG |
| H4K16ac_Rev | GTCGATTATCGCGACGTAA |
| H4K20ac_Rev | GTGATATCGCGTTAACGTCG |
| H3K27acS28ph_Rev | TATCGCGCGAAACGAC |
| tetraAc-H3_(K4/9/14/18ac)_Rev | GTACCGCGCGTATCG |
| tetraAc-H4_(K5/8/12/16ac)_Rev | TCGCATCGCCGAATC |
| tetraAc-H2A_(K5/8/13/15ac)_Rev | TAATCGACGCGTTACGC |
| KLF4_Promoter_For | CACTCGAGAGCGCGATTAT |
| KLF4_Promoter_Rev | CTTCTCCTAGCTTCTGAGATTCC |
| KLF4_gene_body_For | GTAGTGCCTGGTCAGTTCATC |
| KLF4_gene_body_Rev | GCCCGTTCTCTTTCCCTTTA |
